# Changes in Cortical Directional Connectivity during Difficult Listening in Younger and Older Adults

**DOI:** 10.1101/2023.05.19.541500

**Authors:** Behrad Soleimani, I.M. Dushyanthi Karunathilake, Proloy Das, Stefanie E. Kuchinsky, Behtash Babadi, Jonathan Z. Simon

## Abstract

One way to investigate the mechanisms that underlie speech comprehension under difficult listening conditions is via cortical connectivity. The innovative Network Localized Granger Causality (NLGC) framework was applied to magnetoencephalography (MEG) data, obtained from older and younger subjects performing a speech listening task in noisy conditions, in delta and theta frequency bands. Directional connectivity between frontal, temporal, and parietal lobes was analyzed. Both aging- and condition-related changes were found, particularly in theta. In younger adults, as background noise increased, there was a transition from predominantly temporal-to-frontal (bottom-up) connections, to predominantly frontal-to-temporal (top-down). In contrast, older adults showed bidirectional information flow between frontal and temporal cortices even for speech in quiet, not changing substantially with increased noise. Additionally, younger listeners did not show changes in the nature of their cortical links for different listening conditions, whereas older listeners exhibited a switch from predominantly facilitative links to predominantly sharpening, when noise increased.

**Graphical Abstract:** 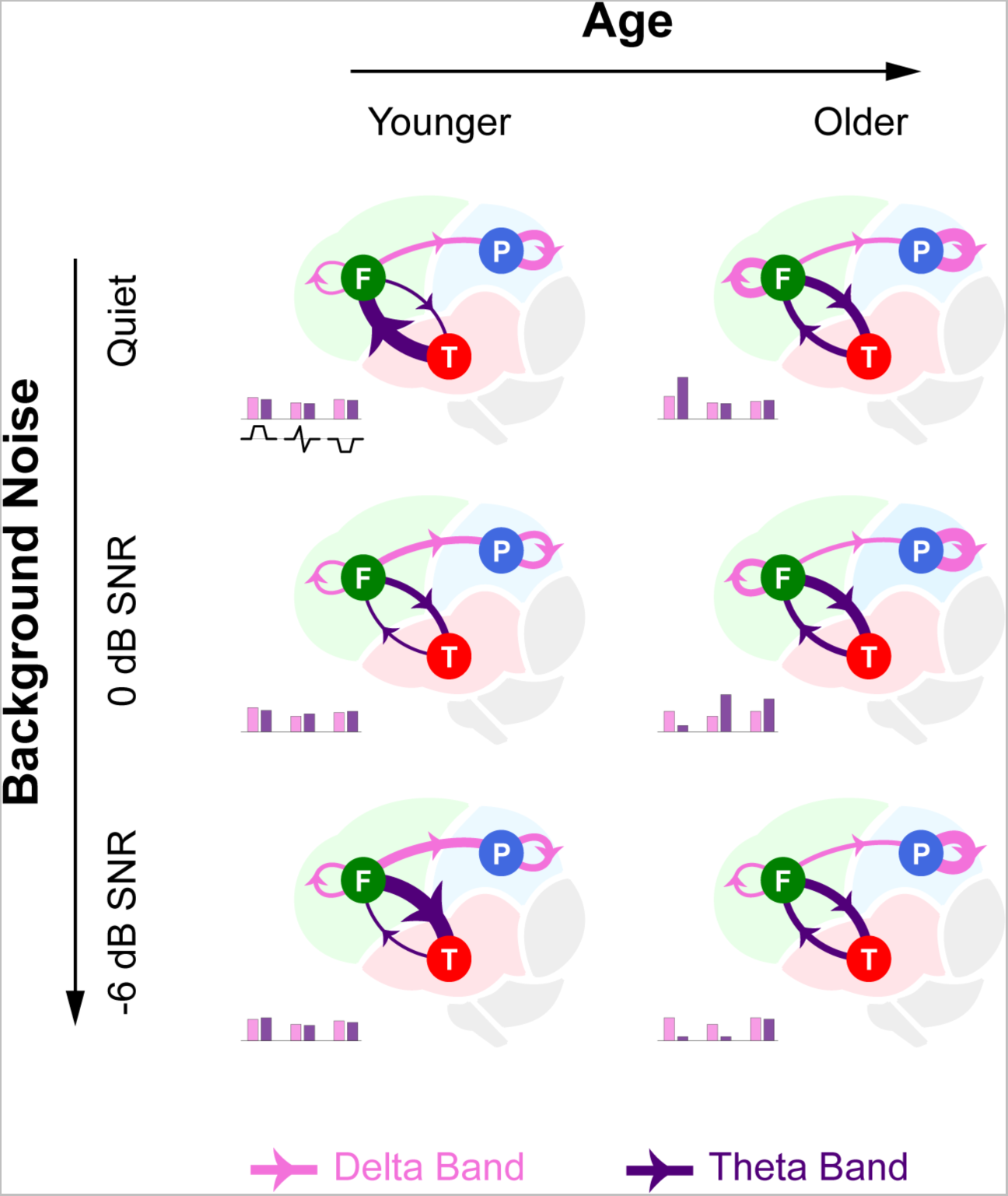

**Highlights:**

- Different bands show strong differences in directional functional connectivity patterns
- Directional functional connectivity patterns altered by listening task difficulty
- Aging dramatically alters directional functional connectivity patterns in during listening
- Nature of functional connectivity, additive vs subtractive, depends on age and task

## INTRODUCTION

The functioning of the neural mechanisms that underlie speech comprehension change with age and under difficult listening conditions (Wong et al., 2009; Karunathilake et al., 2023). Increasing age and listening difficulty has been associated with changes in frontal cortex activity (for a review, see Kuchinsky & Vaden (2020)), which may be indicative of top-down compensatory processing or dedifferentiation, i.e., reduced processing specificity across brain regions (for a review, see Wingfield & Grossman (2006)). Inspecting how cortical connectivity changes with age and difficult listening conditions can provide additional insight into the extent to which neural systems exist to support speech comprehension and the extent to which coordination across regions breaks down with aging (Peelle et al., 2010). Standard functional connectivity studies of aging, in contrast to effective connectivity or other causal modeling approaches, have generally revealed declines in the degree of co-activation of two regions, but cannot speak to changes in directionality (for a review of approaches, see Sala-Llonch et al. (2015)). Insight into directionality may be particularly important when trying to quantify the extent to which age-related changes in connectivity results are driven by changes in bottom-up versus top-down processing. Though functional connectivity is robust and fast to compute (Smith et al., 2013), effective connectivity provides additional insight into directional relationships and are thought to be more biologically meaningful (Friston, 2011).

A well-known *directional* connectivity measure is Granger causality, which assesses the flow of information based on the improvement of temporal predictability of a target signal given the history of a source signal (Bressler & Seth, 2011; Seth et al., 2015). Granger causality has been widely used in functional magnetic resonance Imaging (fMRI) data analysis, and provides insight as to the causal mechanisms of speech and language processing (Upadhyay et al., 2008; Evans & McGettigan, 2017). For instance, such existing evidence supports the causal contribution of motor and visual cortices during speech perception tasks (Van Atteveldt et al., 2009; Osnes et al., 2011; Behroozmand et al., 2015), and illustrates age-dependent cortical connectivity patterns in the auditory cortex (Fei et al., 2020; Rysop et al., 2022).

fMRI has relatively high spatial resolution, but is only able to capture slow neural interactions (typically ~1 Hz), not those at higher frequencies known to be critical for complex speech and language processing (Bastiaansen et al., 2005; Riecke et al., 2009; Ding & Simon, 2012; Mai et al., 2016; Pu et al., 2020; Leicht et al., 2021). Magnetoencephalography (MEG) is another non-invasive neuroimaging technique that achieves higher temporal resolution, or the order of milliseconds (Baillet et al., 2001). As opposed to fMRI, however, MEG only provides indirect measurements of the underlying neural sources (Baillet et al., 2001). As such, to extract the cortical level connectivity from MEG recordings, two stages typically needed: first, the neural source activities are estimated via standard source-localization methods, and then the desired connectivity analysis is carried out on the estimated source time courses (Schoffelen & Gross, 2009). However, due to the ill-posed nature of the MEG source-localization problem, any statistical biases arising as part of solving the source localization problem, e.g., localization spread, automatically propagate to the subsequent connectivity analysis stage, resulting in an abnormally high false alarm rate (Palva et al., 2018; Soleimani et al., 2022).

Instead, here we utilized a recently introduced methodology for MEG directional functional connectivity analysis called network localized Granger causality (NLGC), which reliably captures Granger causal interactions in a single-stage analysis, without requiring any intermediate source-localization step (Soleimani et al., 2022). As an immediate advantage, NLGC allows the use of MEG recordings to measure directional functional connectivity throughout the entire cortex with high statistical precision, a feat previously requiring either (slow) fMRI or invasive recordings in animal models (Rauschecker & Scott, 2009; Wang et al., 2012; Peelle, 2016). Secondly, in this work, we leveraged NLGC’s incorporation of sparse vector autoregressive models to quantify the nature of the inferred directional links in terms of the facilitative (additive), suppressive (subtractive), or sharpening (serial additive/subtractive) contribution of the source to target. This allowed comparison of, and extensions to, existing results showing distinct roles for excitation and inhibition, e.g., excitation/inhibition balance in the auditory cortex modulated by speech demand in older adults (Dobri & Ross, 2021).

We applied NLGC to a set of experimentally recorded MEG data from an auditory task where younger and older participants listened to 1-minute-long speech segments under varying background noise levels that altered listening difficulty. We focused on the directional cortical connectivity patterns in the frontal, temporal, and parietal lobes within the delta (0.1-4 Hz) and theta (4-8 Hz) frequency bands. We investigated the impact of age on this auditory network connectivity, examining changes in the strength of connections within the auditory areas across different conditions. Our findings revealed distinct age-, frequency- and condition-related changes in the connectivity. Specifically in the theta band, as the speech listening task was made more difficult, we showed increased frontal-to-temporal connections and correspondingly decreased temporal-to-frontal connections in younger adults. In contrast, in older adults, we observed bidirectional connectivity between frontal and temporal cortices even for speech in quiet, suggesting that auditory speech processing even of non-degraded speech requires additional resources from the frontal cortex. Additionally, younger listeners did not show any changes in the nature of their cortical links for different listening tasks, whereas in older listener there was a switch from predominantly facilitative links to dominantly sharpening, when the task changed from listening to speech in quiet to listening to noisy speech. The delta band, in contrast, showed no significant changes in the distribution of links across age, condition, or in the nature of the links.

## RESULTS AND DISCUSSION

### NLGC Reveals Source-level Directional Connectivity Patterns from Sensor-level MEG Data

To quantify the Granger causal link from brain region R_1_ to region R_2_, one defines two competing models, the *full* and *reduced* models. The full model includes all other (not just R_1_ and R_2_ but including them) sources’ past activity in predicting the activity of R_2_, whereas in the reduced model the past activity of R_1_ is omitted as a predictor. If the prediction error of the reduced model is significantly larger than that of the full model, which indicates that the removing knowledge of the past of R_1_ significantly degraded the prediction of R_2_, then we say there is a significant Granger causal (GC) link from R_1_ to R_2_. NLGC performs this procedure for all source-target pairs in the source space, though directly using the sensor-level MEG data, ultimately providing a source-level directional connectivity map (see Methods).

As a first illustrative example, we examined the directional connectivity in the theta band (4-8 Hz) between the frontal and temporal regions of the brain under three different listening conditions (speech in quiet, and in background noise at two different signal-to-noise ratios (SNR): 0 and −6 dB) for two different age groups (younger versus older adults). Fig. 1A shows the frontal-to-temporal and temporal-to-frontal connections (GC links) for both the grand average across individuals and for representative individuals across age groups and listening conditions. In younger adults, as the speech listening task is made more difficult, we saw increased frontal-to-temporal connections (red arrows) and correspondingly decreased temporal-to-frontal connections (green arrows), whereas in older adults, we show bidirectional connectivity between frontal and temporal cortices even for speech in quiet.

**Figure 1.**
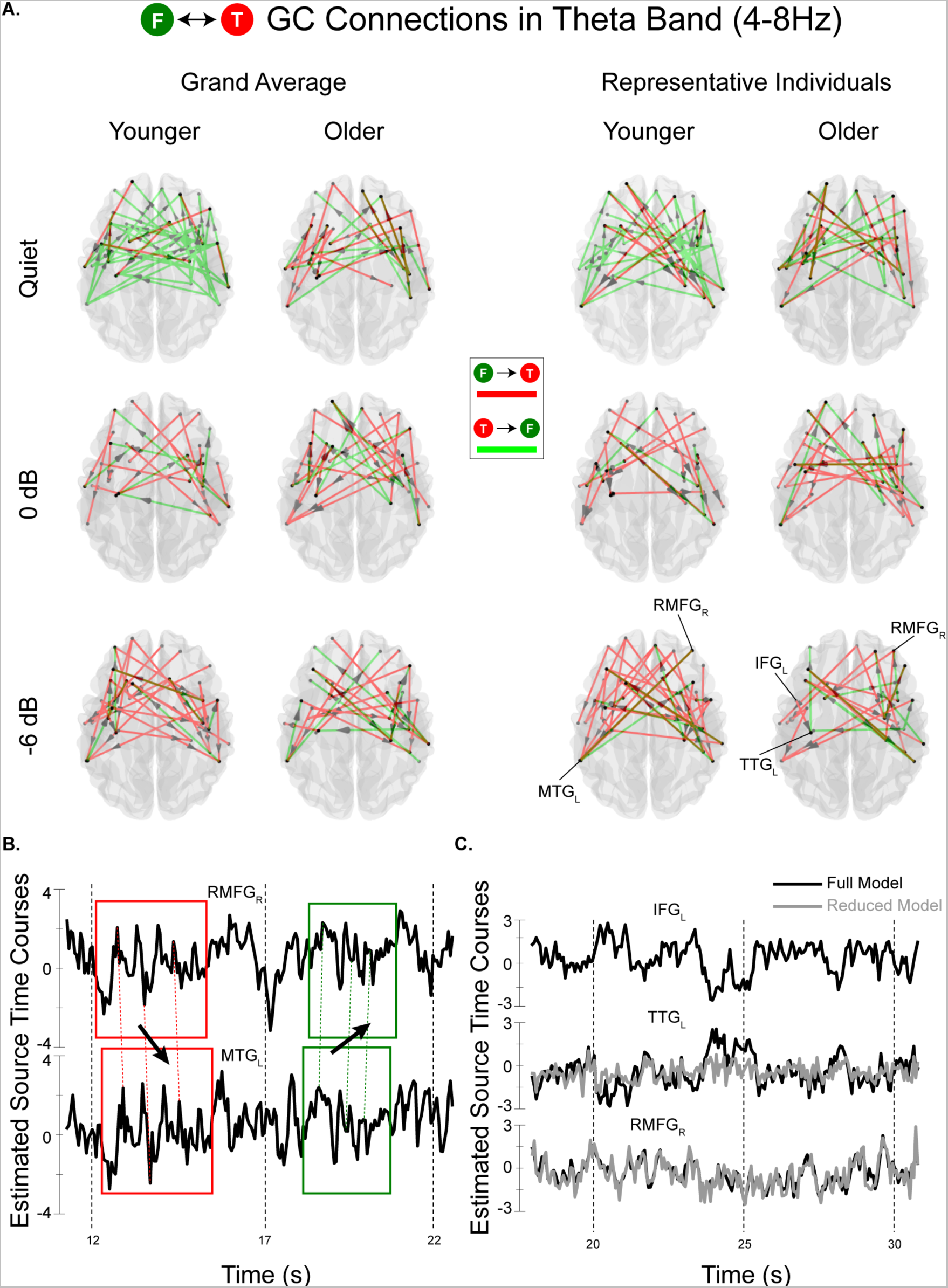
Age-related changes in directional connectivity in the theta band (4-8 Hz) between frontal and temporal regions under different listening conditions. **A.** frontal-to-temporal (red) and temporal-to-frontal (green) connections for the group averages (left) and representative individuals (right) across listening conditions and age group. **B.** Average activity of the ROIs RMFG_R_ (right rostral middle frontal gyrus) and MTG_L_ (left rostral middle frontal gyrus), illustrating a bidirectional GC link. **C.** The activity of three ROIs IFG_L_ (left inferior frontal gyrus), TTG_L_ (left transverse temporal gyrus), RMFG_R_ and their corresponding reduced models of the GC link IFG_L_-to-TTG_L_ and a non-GC link from IFG_L_ to RMFG_L_.

Bidirectional GC links can be best understood by inspecting the time-course of source activities underlying a few representative GC links. Fig. 1B shows the average activity of two representative sources: the right rostral middle frontal gyrus (RMFG_R_) and the left middle temporal gyrus (MTG_L_), which are found to have *bidirectional* Granger causal links. Expectedly, we observed that in the first epoch (marked by red boxes) the activity pattern of RMGF_R_ precedingly appears in MTG_L_, and in the second epoch (marked by green boxes), the activity of MTG_L_ leads that of RMFG_R_. NLGC was indeed able to capture this bidirectional statistical relationship between the two time series, which is in general distinct from the bidirectional links obtained by correlation-based methods to quantify functional connectivity.

As an addition illustrative comparison, Fig. 1C shows the source time-courses underlying the GC links between left inferior frontal gyrus (IFG_L_), left transverse temporal gyrus (TTG_L_), and RMFG_R_. Here, we inspect two possible GC links: IFG_L_-to-TTG_L_, which was found to be significant by NLGC, and IFG_L_-to-RMFG_R_, which was *not* found to be significant. The first row shows the activity of IFG_L_ as the source. The second row shows the predicted activity of TTG_L_ using the full model (black trace) and the reduced model (gray trace), in which the activity of IFG_L_ is removed as a predictor. As it can be observed, the two predictions were quite distinct, which is consistent with a GC link from IFG_L_ to TTG_L_. On the other hand, the full and reduced model predictions of RMFG_R_ shown in the third row were nearly identical, which is consistent with no GC link from IFG_L_ to RMFG_R_.

### Age-related Top-Down and Bottom-Up Changes in the Theta Band

Fig. 1A suggests that there were age-related changes in the directionality and strength of connectivity in the theta band between the frontal and temporal regions of the brain under difficult listening conditions, which could be critical for understanding age-related changes in speech processing and cognition (Wong et al., 2010; Hipp et al., 2011; Peelle, 2016).

To investigate this finding more fully, we inspected theta band connectivity between temporal, frontal, and parietal regions quantitatively. Fig. 2A shows a graphical representation of Fig. 1A (left column), where the average percentage of GC links for each age group and listening condition are shown by arrows connecting cortical regions (frontal = green, parietal = blue, and temporal = red); the grayscale levels represent the percentage of links for each connectivity pair. Average activity power corresponding to each region is also indicated by correspondingly color-coded gauges. As can be seen, younger listeners showed predominantly temporal-to-frontal connectivity in the speech-in-quiet condition, while in noisy conditions, the pattern was reversed to frontal-to-temporal connectivity. Older listeners, on the other hand, displayed predominantly frontal-to-temporal connectivity regardless of the listening condition. Additionally, in the 0 dB SNR condition, the majority of parietotemporal links were from temporal to parietal for younger listeners, whereas in older listeners, the pattern was reversed.

**Figure 2.**
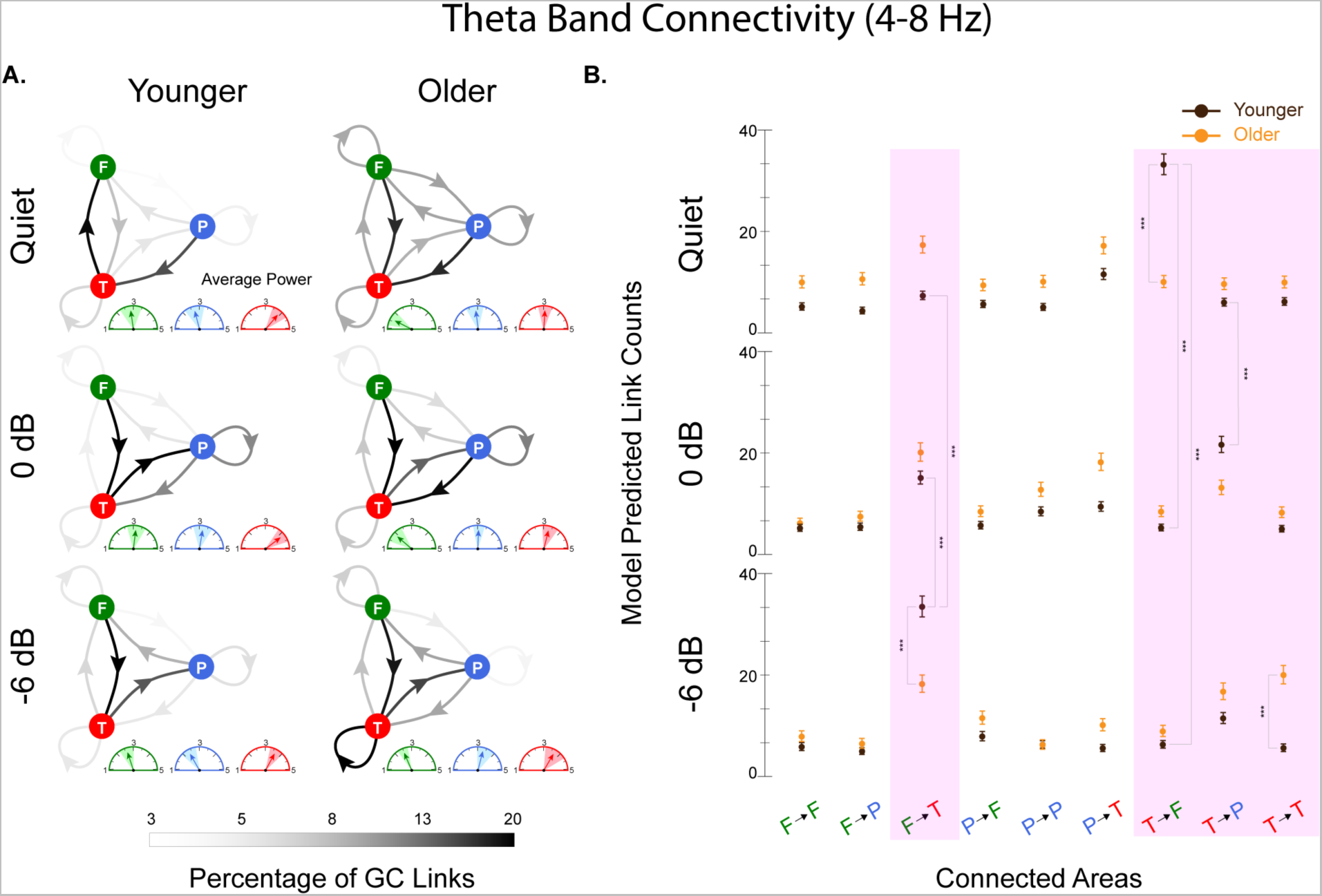
Theta band connectivity across age and listening condition. **A.** Group average percentages of GC links in and across three regions of interest (ROI)s: frontal (F) = green, parietal (P) = blue, temporal (T) = red. Grayscale level represents the connection strength at the group average level for each age group and listening condition. Group average power corresponding to each ROI is shown by colored gauges, where color shades around the arrows show 95% confidence interval of the distribution of the average power across participants. **B.** Statistical model results for the number of links per connectivity region pair for younger (brown) and older (orange) adults per listening condition. Significant differences are highlighted in pink (*** *p* < 0.001, ** *p* < 0.01, * *p* < 0.05)

Additionally, average power in all regions generally increased with background noise for older adults, whereas for younger listeners, the average power change did not generally follow listening difficulty. In general, high power in a source does not generally indicate a strong GC influence on the target (Gross et al., 2001), which has been previously noted in studies that have reported weak or absent correlations between Granger causality and measures of neural activity such as power or coherence (Seth, 2007; Dhamala et al., 2008). These findings suggest that while activity power and Granger causality are related measures of neural activity, they capture different aspects of neural dynamics and can be interpreted independently.

In order to investigate all interacting effects of age group, listening condition, and connected areas, we used a generalized linear mixed-effects model to predict the number of links per connectivity regions for younger and older participants and per listening condition in the theta band (see Supplemental Table S1). The results displayed in Fig. 2B reveal statistically significant differences in temporal-to-frontal and frontal-to-temporal connectivity, as well as in temporal-to-parietal and temporal-to-temporal connectivity. Fig. 3 shows a graphical representation of connectivity differences across age and listening conditions (the connectivity arrows are filled with “+” and “-“ signs, reflecting increase and decrease in connectivity, respectively). In Fig. 3A, age-group differences are shown per listening condition; Fig. 3B illustrates the changes across listening conditions for each age group. Differences in average power are also shown. All links were statistically significant, but not all differences were necessarily of large effect size; pink daggers mark significant differences with a substantial effect size of at least d = 30, the 95% quantile of effect size. The results with the largest effect sizes can be summarized as follows:

1. For younger adults, the decrease in temporal-to-frontal connectivity for speech in quiet compared to the 0 dB SNR and −6 dB SNR conditions, and the corresponding increase in frontal-to-temporal activity for speech in quiet and 0 dB SNR conditions to −6 dB SNR, are not only significant but also of large effect size. For older adults, the bidirectional connectivity pattern shows significant changes with changes in noise condition but not consistently and with small effect size. Additionally, there was significantly less, and with large effect size, temporal-to-frontal connectivity during speech-in-quiet listening in older adults than younger, and there was significantly less, and with large effect size, frontal-to-temporal connectivity during −6 dB SNR listening in older adults than younger. The reversal of the frontotemporal connectivity direction observed in younger and older adults with the listening difficulty (Fig. 3A, top and bottom graphs) is consistent with previous work. For example, a study by Obleser et al. (2007) found that in a listening task with competing speakers, younger adults showed increased functional connectivity between temporal and frontal regions, while older adults showed decreased connectivity between these regions. Additionally, a study by Peelle (2013) found that during speech comprehension, younger adults showed greater theta-band coherence between frontal and temporal regions, while older adults showed reduced coherence between these regions.
2. The relationship between neural activity power and the underlying cognitive processes is complex. As such, there was no direct relationship between the neural source power and Granger causality, as is reflected by the power difference gauges in Fig. 3. However, it has been suggested that the increase in power may reflect compensatory neural mechanisms in response to cognitive decline in aging (Klimesch, 1999). In terms of the effect of background noise on neural activity power, a study by (Anderson et al., 2012) found that in a speech perception task, younger adults showed a reduction in theta-band power in with respect to increased background noise, while older adults showed an increase in power.
3. In the most difficult listening condition (−6 dB SNR), older adults show significantly enhanced temporal-to-temporal connectivity, with large effect size. The increased temporal-to-temporal connectivity in older adults in the −6 dB SNR condition (Fig. 3A, bottom graph) is consistent with a study by (Wostmann et al., 2016), which found that in a speech-in-noise task, older adults showed increased connectivity within temporal regions in the theta band compared to younger adults. The authors suggested that this increase in connectivity may reflect compensatory mechanisms in response to age-related decline in auditory processing.
4. Younger adults showed significantly enhanced temporal-to-parietal connectivity, with large effect size, when switching from speech in quiet to the two noisy conditions, and significantly decreased parietal-to-temporal connectivity as well, though with a smaller effect size. Older adults also showed both these patterns but with a smaller effect size. Though previous fMRI studies have noted the role of a temporal-parietal system for semantic memory for external stimuli (Binder et al., 2009), to our knowledge, the pattern observed here has not been reported previously.
5. Older adults show significantly greater frontal-to-frontal connectivity and greater bidirectional frontal-to-parietal connectivity than younger adults for all noise conditions, though with a smaller effect size. To our knowledge, this pattern has not been previously observed in the directed connectivity literature.

**Figure 3.**
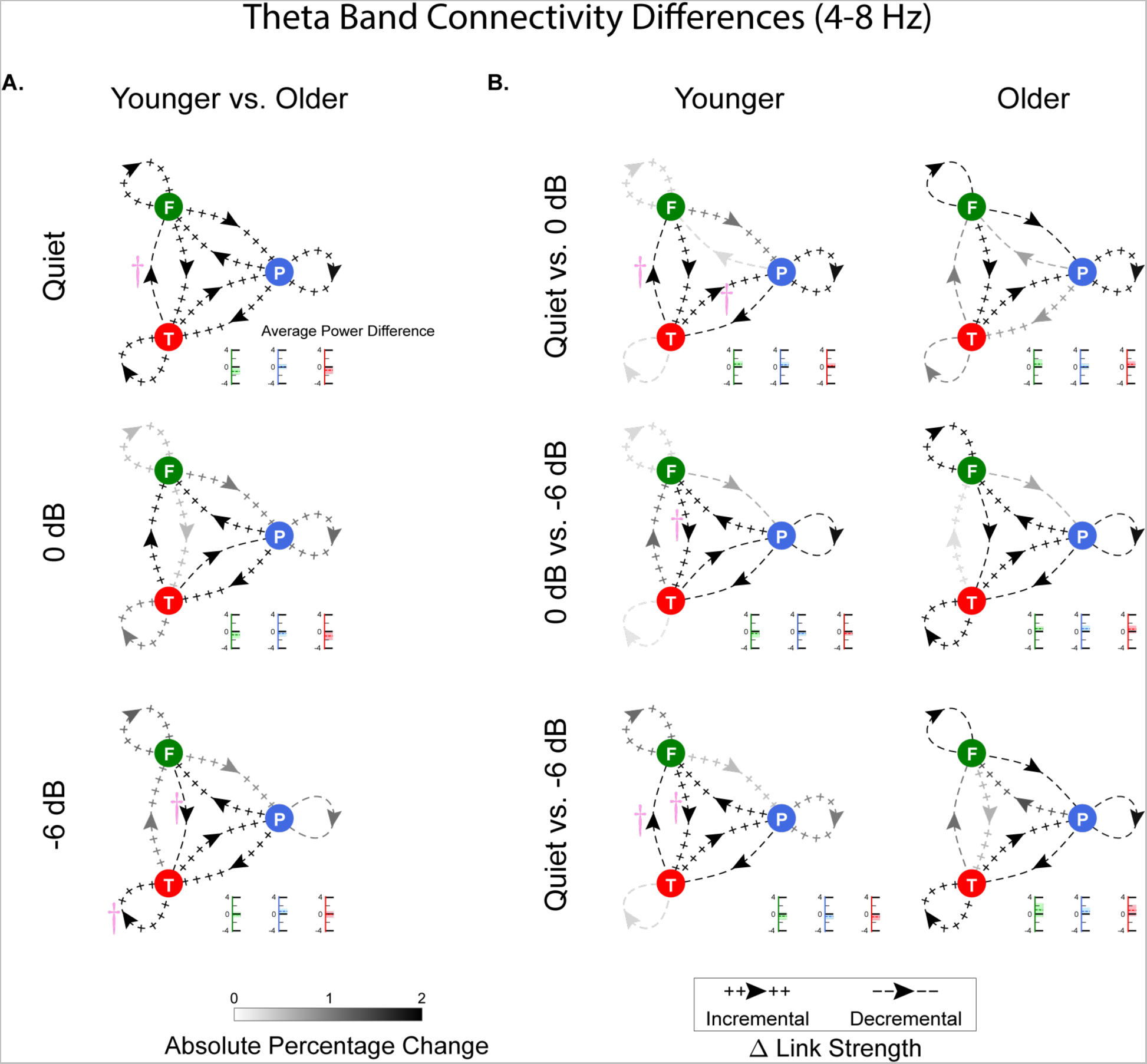
Age- and condition-related connectivity differences in theta band. **A.** Grand average percentage changes from younger to older adults (frontal (F) = green, parietal (P) = blue, temporal (T) = red), across listening conditions. The grayscale level and positive/negative signs of the arrows show the absolute change and its sign, respectively. Grand average power changes corresponding to each ROI are shown by the correspondingly colored gauges, where colored shades around the arrows show 95% confidence intervals across participants. B. The analogous noise-condition-related group average of within-subject changes for both age groups. All displayed connectivity changes are significant; pink daggers indicate connectivity changes with effect size *d* at the 95% quantile and above.

We investigated, but did not observe, any statistically significant changes across hemispheres. In summary, these results suggest that aging and listening difficulty interactively affect the brain’s connectivity. Directional connectivity analyses help to advance our understanding of age-related changes in speech processing by highlighting potentially causal, compensatory mechanisms that may be employed by the older brain to adapt to challenging listening environments (i.e., changes in bottom-up temporal cortex activity signaling top-down frontal systems) and overcome age-related dedifferentiation of neural processing specificity (Wingfield & Grossman (2006).

### Age-Related Reversal in Frontal-Parietal Connectivity in the Delta band

We similarly investigated directional connectivity in the delta band (0.1-4 Hz). Fig. 4 shows the corresponding links between frontal and parietal areas overlaid on brain plots, analogous to the plots shown in Fig. 1A. While the number of frontal-to-parietal GC links (blue arrows) and parietal-to-frontal GC links (green arrows) are more balanced for younger adults for all noise conditions, it can be seen that older adults exhibit more parietal-to-frontal GC links. Interestingly, this age-related change in the directionality will be seen to be independent of the difficult listening condition. This result underscores the importance of considering directionality in connectivity analysis. For instance, authors in (Eckert et al., 2012; Peelle, 2016) showed that in older adults the correlation between parietal and frontal areas are increased compared to the younger participants. While our results are consistent with existing work, it can potentially complement such findings by highlighting the underlying age-related changes in the directionality of auditory information flow (Anderson et al., 2010; Geerligs et al., 2015). Fig. 5A shows the graphical representation of delta band connectivity across age and listening conditions, (in the same format as Fig. 2). The overall pattern of connectivity is quite different from that seen in the theta band, e.g., a more prominent frontoparietal connectivity in delta compared to theta (Fig. 2A).

**Figure 4.**
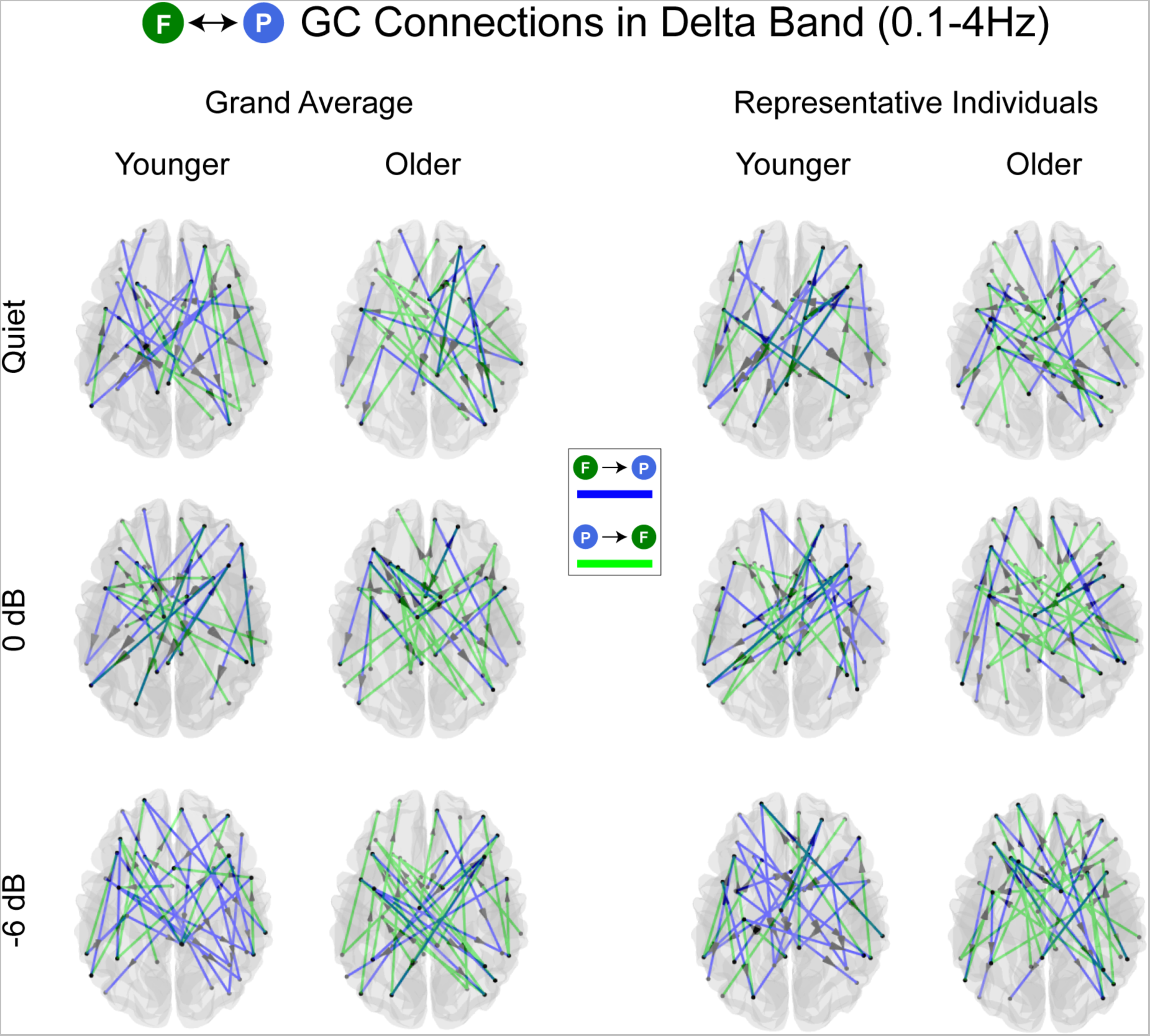
Age-related changes in directional connectivity in the delta band (0.1-4 Hz) between frontal and parietal lobes under different listening conditions. frontal-to-parietal (blue) and parietal-to-frontal (green) connections for the group averages (left) and representative individuals (right) across listening conditions and age group.

**Figure 5.**
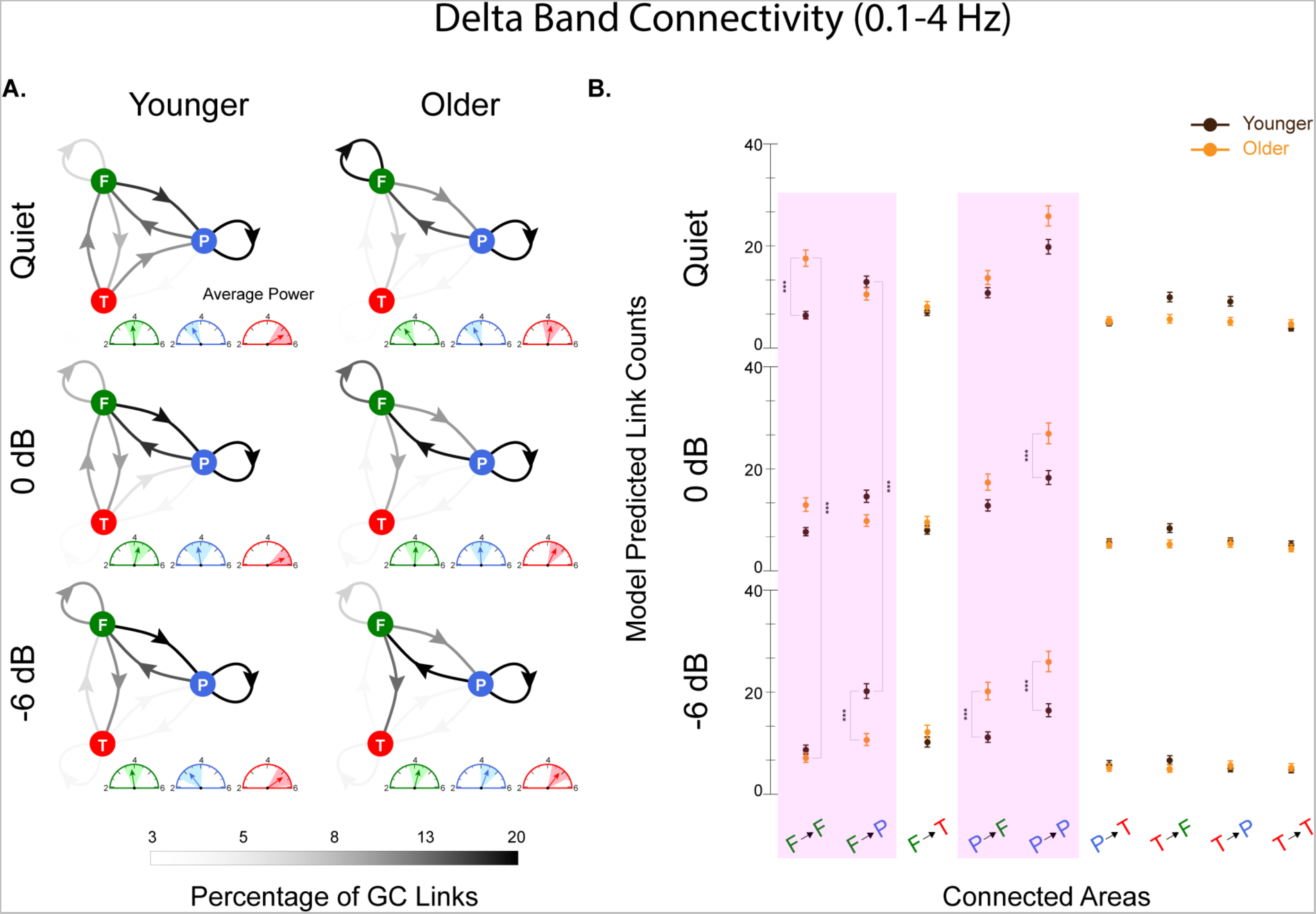
Delta band connectivity across age and listening condition. **A.** Group average percentages of GC links in and across three regions of interest (ROI)s: frontal (F) = green, parietal (P) = blue, temporal (T) = red. Grayscale level represents the connection strength at the group average level for each age group and listening condition. Group average power corresponding to each ROI is shown by colored gauges, where color shades around the arrows show 95% confidence interval of the distribution of the average power across participants. **B.** Statistical model results for the number of links per connectivity region pair for younger (brown) and older (orange) adults per listening condition. Significant differences are highlighted in pink (*** *p* < 0.001, ** *p* < 0.01, * *p* < 0.05)

In order to investigate all interacting effects of age group, listening condition, and connected areas, we used another generalized linear mixed-effects model to predict the number of links per connectivity regions for younger and older participants and per listening condition in the delta band (Supplemental Table S1). The results displayed in Fig. 5B reveal statistically significant differences in parietal-to-frontal and frontal-to-parietal connectivity, as well as in parietal-to-parietal and frontal-to-frontal connectivity. In Fig. 6A, age-group differences are shown per listening condition; Fig. 6B illustrates the changes across listening conditions for each age group. Differences in average power are also shown. All links are statistically significant, but not all differences are necessarily of large effect size; pink daggers mark significant differences with a substantial effect size of at least d = 15, the 95% quantile of effect size. The results with the largest effect sizes can be summarized as follows:

1. Older adults exhibited significantly more frontal-to-frontal connections, and with large effect size, compared to younger adults in the speech in quiet condition only (Fig. 6A). Consistent with this finding, Alain & McDonald (2007) found that older adults exhibit more correlated activity between areas within frontal cortex compared to younger adults.
2. For both the 0 dB SNR and −6 dB SNR noise conditions, there were significantly more parietal-to-parietal connections, and with large effect size, in older compared to younger adults. To our knowledge, this has not been previously observed in the literature.
3. In the most difficult listening condition, −6 dB SNR, older adults exhibited significantly increased parietal-to-frontal and decreased frontal-to-parietal connections compared to younger adults, both with large effect size. Although previous studies have observed age-related increases in fronto-parietal compensatory recruitment (e.g., Sharp et al. (2006)), this also has not been studied before in terms of the directionality and potentially causal interactions between these regions.
4. Frontal-to-parietal connections exhibited an significantly increasing trend with listening difficulty in younger adults, with large effect size, consistent with an existing study (Park et al., 2015) showing coherence changes between frontal and parietal at low frequencies.
5. Frontal-to-frontal connectivity significantly decreased in older adults as the noise level increased, with large effect size. This was accompanied with a significant overall decrease in the connectivity between frontal and temporal regions, with smaller effect size. This is consistent with the findings of (Peelle, 2016), who point out evidence from existing literature that older adults with hearing loss exhibit decreased functional connectivity between frontal and temporal regions. This disruption in connectivity is posed as being related to the difficulty of speech processing in noisy environments for older adults.

**Figure 6.**
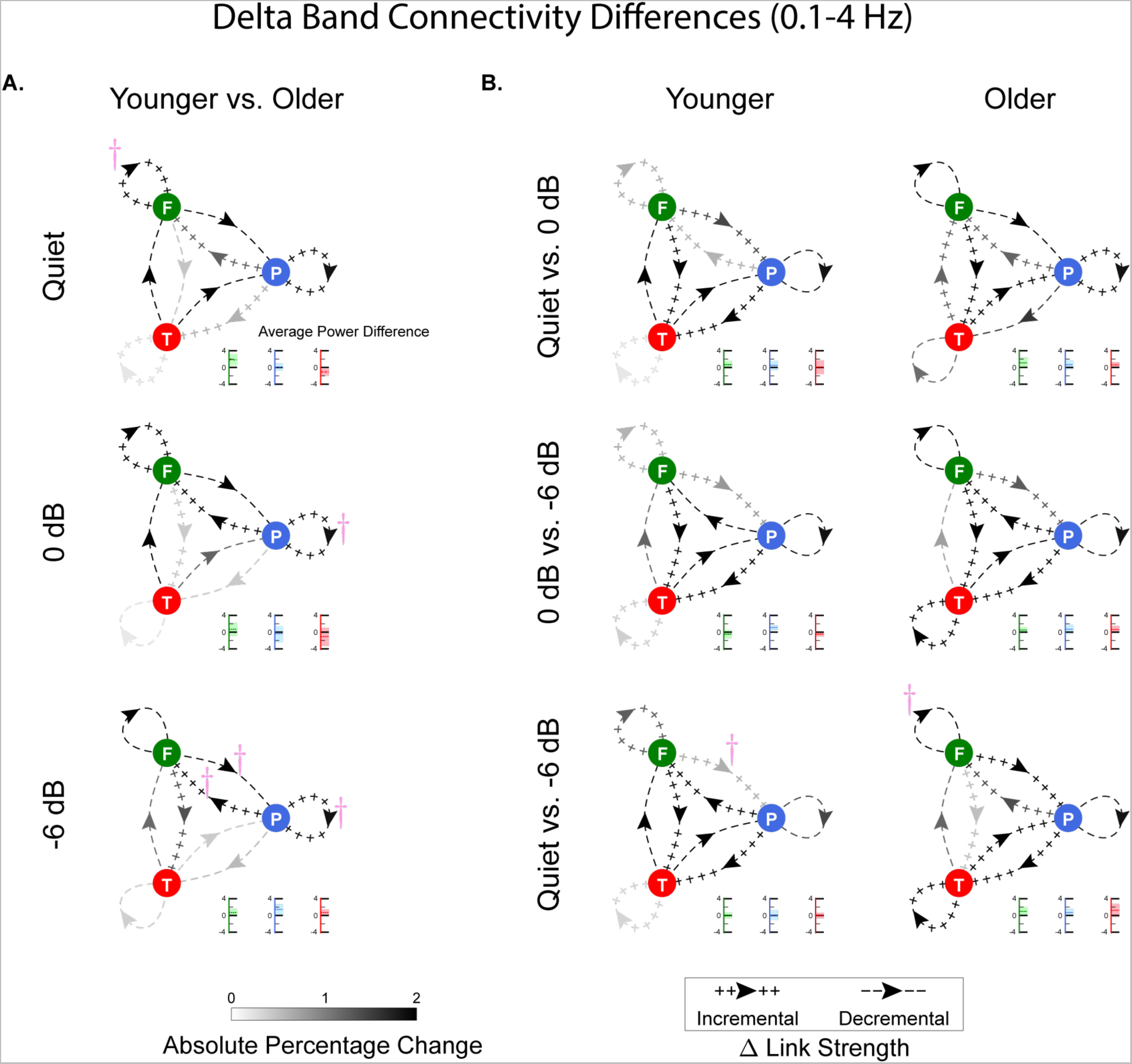
Age- and condition-related connectivity differences in delta band. **A.** Grand average percentage changes from younger to older adults (frontal (F) = green, parietal (P) = blue, temporal (T) = red), across listening conditions. The grayscale level and positive/negative signs of the arrows show the absolute change and its sign, respectively. Grand average power changes corresponding to each ROI are shown by the correspondingly colored gauges, where colored shades around the arrows show 95% confidence intervals across participants. B. The analogous noise-condition-related group average of within-subject changes for both age groups. All displayed connectivity changes are significant; pink daggers indicate connectivity changes with effect size *d* at the 95% quantile and above.

In summary, observed age-related increases in within-lobe connectivity (i.e., frontal-to-frontal and parietal-to-parietal) and decreases in across-lobe connectivity (i.e., weaker temporal-parietal effects) in at least some listening conditions appear to align with the finding that aging is associated with increases in short-range connections and decreases in long-range connections (Tomasi & Volkow, 2012).

### Nature of Links Varies Across Age and Condition Only for Theta Band

Another outcome of the NLGC framework is the ability to characterize GC links in terms of the nature of the interactions between the source-target pair (See Methods and Supplementary Table S2). Fig. 7 illustrates these results in the theta band. For younger adults, no significant differences were seen across noise conditions or connectivity nature, in strong contrast to the many significant differences seen for older adults. Across age groups, connectivity significantly increased in older adults for facilitative connections only for speech in quiet, for sharpening connections in all noise conditions, and not at all for suppressive connections. Additionally, connectivity significantly decreased in older adults for facilitative connections only in the 0 dB SNR conditions, not at all for the sharpening connections, and for suppressive connections in both the speech in quiet and 0 dB SNR conditions. The general decrease in suppressive connectivity is consistent with the aging-related decrease in GABA-based inhibition seen in many auditory studies, both animal (Willott et al., 1991; Caspary et al., 1995; Hughes et al., 2010; Caspary et al., 2013; Richardson et al., 2013; Parthasarathy et al., 2019; Ramamurthy & Recanzone, 2020) and human (Lalwani et al., 2019; Ross et al., 2020; Dobri & Ross, 2021; Harris et al., 2022). More interesting, perhaps, is the dynamic tradeoff between facilitative and sharpening connectivity seen in older adults when shifting from listening to speech in quiet (maximal facilitative and minimal sharpening) to speech in noise (reduced facilitative but increased sharpening). It may ultimately be possible to equate facilitative connectivity with dominantly excitatory connections, suppressive connectivity with dominantly inhibitory connections, and sharpening connectivity with a temporally delayed mix of the two. If this were true, it would correspond to older adults exhibiting an expected imbalance of excitatory and inhibitory connections while listening to speech in quiet, but that in the presence of noise there occurring a functional shift of a substantial number of excitatory connections from purely facilitative (excitatory connections only) to sharpening (a temporally mix of excitatory and inhibitory connections). The findings here generally support existing studies that back the hypothesis of excitatory/inhibitory balance modulation by the acoustic features of speech and speech processing demand (Kuchibhotla & Bathellier, 2018; Kondo & Lin, 2020).

**Figure 7.**
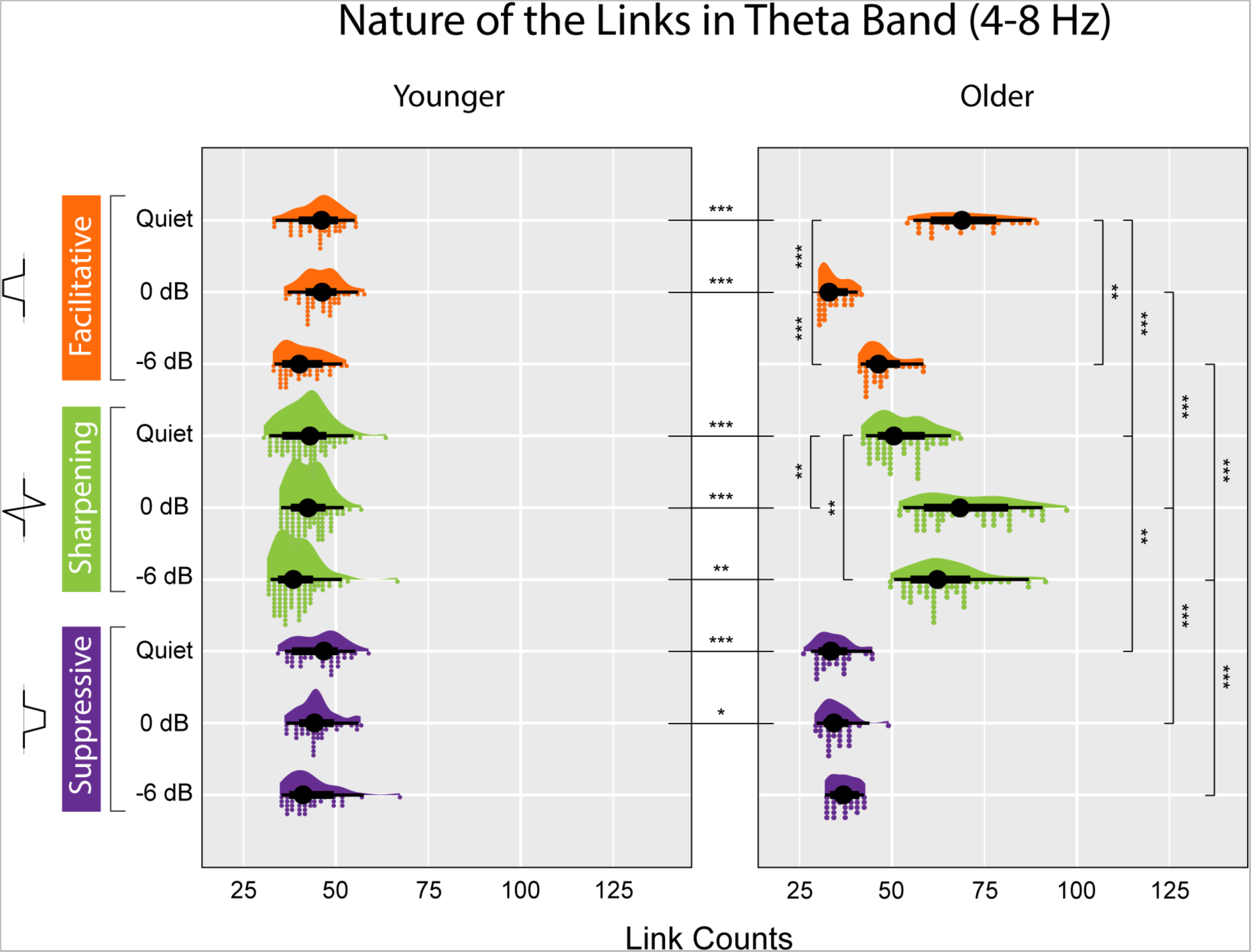
Nature of the link analysis in theta band per age and listening condition. Left column: younger adults, right column: older adults. Three types of the links are shown: facilitative (dark orange), sharpening (light green), and suppressive (dark purple). Significant differences (not seen within younger adults) were obtained via post hoc testing are marked according to their corresponding Holm-corrected *p*-value (*** *p* < 0.001, ** *p* < 0.01, * *p* < 0.05).

The same analysis performed over the delta band (Table S2), however, as illustrated in Fig. 8, showed no statistically significant differences within or across age groups. Additional statistical analysis in both bands showed no dependence of the distribution of the nature of the links, as shown in Figs. 7 and 8, with connectivity regions, e.g., temporal-to-parietal or frontal-to-temporal, for either of the frequency bands.

**Figure 8.**
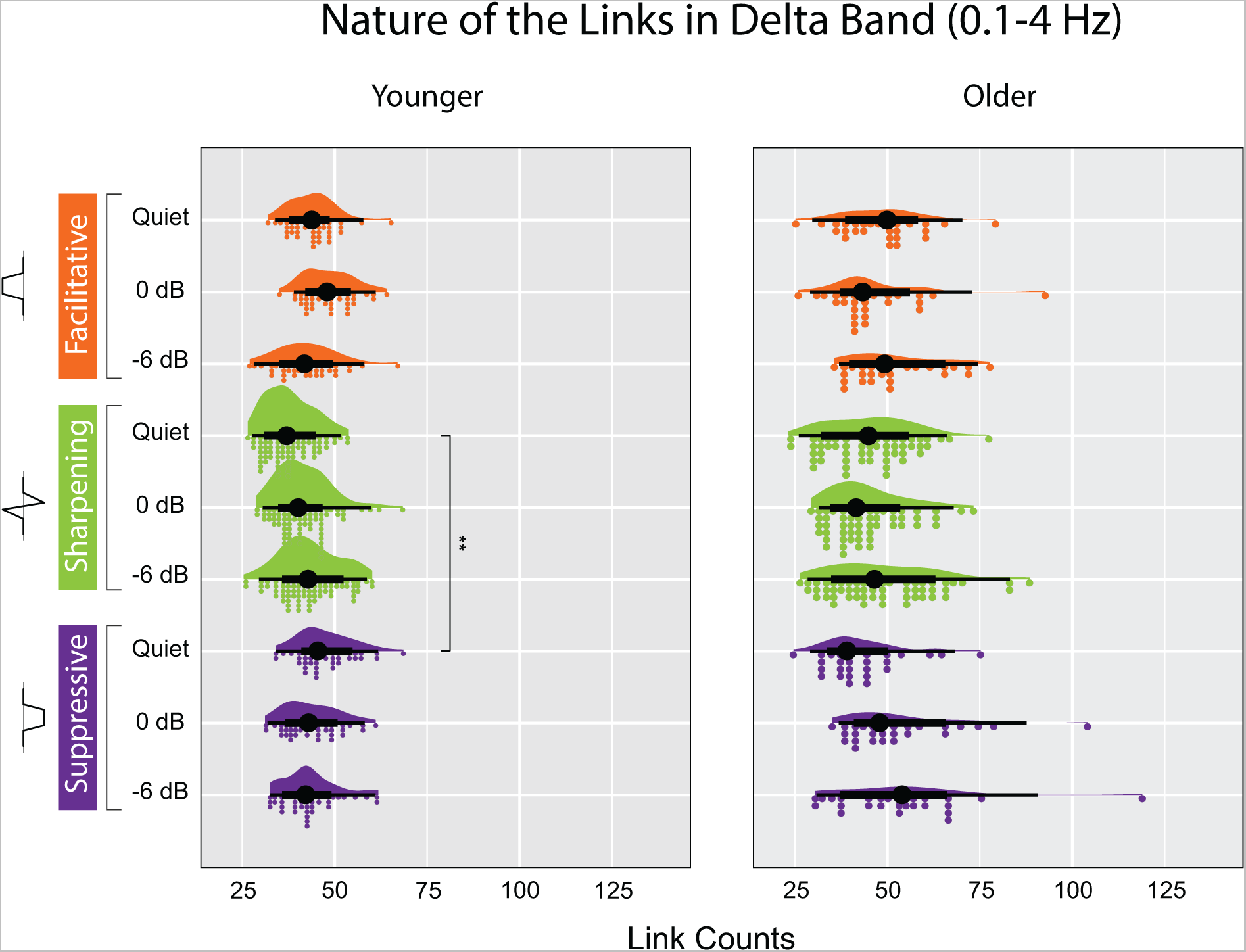
Nature of the link analysis in delta band per age and listening condition. Left column: younger adults, right column: older adults. Three types of the links are shown: facilitative (dark orange), sharpening (light green), and suppressive (dark purple). Significant differences (not seen within older adults, one seen within younger adults) were obtained via post hoc testing are marked according to their corresponding Holm-corrected *p*-value.

### Benefits provided by NLGC

This study provides novel insights into network connectivity and its changes that differ from previous human non-invasive neurophysiological studies in several key aspects:

- **Reliable network localization, as opposed to source localization:** NLGC offers a robust and reliable solution for obtaining directional functional connectivity without the need for intermediate source localization, which is a step commonly used in two-stage methods that identify connectivity changes in the auditory cortex (Paraskevopoulos et al., 2017; Paraskevopoulos et al., 2019; Tait & Zhang, 2022). This streamlining of the analysis pipeline, namely network localization, reduces potential errors and increases the efficiency of the analysis (Palva & Palva, 2012).
- **Identifying directionality**: Our analysis provides directionality of the functional connections, which could point to the flow of information. Undirected metrics used in previous studies, such as Pearson correlation coefficient (Vanneste & De Ridder, 2015; Zhang et al., 2016), may not capture this important aspect of neural communication.
- **Capturing nature of connections**: NLGC allows to categorize the nature of the captured relationships as facilitative, suppressive, or sharpening. This categorization can be thought of as the meso-scale equivalent of the excitation/inhibition relations at the neuronal level (Pantev et al., 2004; Pantev, 2012; Stein et al., 2013; Inui et al., 2016; Tada et al., 2020) and provides a novel perspective on the network-level relationship between the activity of different ROIs.

Although NLGC offers many advantages in terms of network connectivity analysis over standard functional connectivity methods, it also has limitations. First, it is more computationally demanding than the two-stage procedures, which can make hyperparameter tuning time-consuming. For this reason, the connectivity regions were restricted to entire cortical lobes rather than a finer parcellation (in turn limiting the ability to distinguish, e.g., parietal contributions to the frontoparietal network from parietal contributions to the default mode network). In addition, the algorithm is sensitive to external artifacts in the neural recording, which may accordingly require additional pre-processing and data cleaning to ensure the accuracy and consistency of the results. These challenges are discussed further in Soleimani et al. (2022).

### Summary

This study provides novel insights into network connectivity important for speech processing at several levels. First, the connectivity in two different bands and across brain areas was shown to depend not only connectivity areas and age group but also on the direction of the connection, sometimes dramatically, and analysis was unaffected even when connectivity was strongly bidirectional. Specifically, the connectivity patterns seen here for the temporal and frontal lobes in the theta band, and for the parietal and frontal lobes in the delta band, confirmed previous results based on auditory aging and difficult listening conditions, but also provided novel results unseen using fMRI or traditional functional connectivity methods. Secondly, the analysis of the nature of the connectivity (facilitative, sharpening, and suppressive) provided novel results showing evidence for connectivity dynamics not just in location but in kind, even hinting at reallocation of excitatory and inhibitory connections at macroscopic scales.

In summary, employing NLGC to study age-related changes in the brain provides a network-level cortical functional connectivity account of auditory processing as a function of frequency, age, spatial distribution of the connections, and listening difficulty.

## METHODS

### Subjects, Auditory Tasks, and Recordings

The experimental dataset used for this study was a part of previous study described in detail in (Karunathilake et al., 2023). Out of 36 total participants who completed the MEG experiment, the current analysis included 22 participants, 13 younger adults (5 males; mean age 21.1 years, range 17–26 years) and 9 older adults (3 males; mean age 69.6 years, range 66-78 years) who completed the structural MRI scans. Additionally, 2 subjects were excluded due to bad fiducials measurements. All participants had normal hearing (125–4000 Hz, thresholds ≤ 25 dB hearing level, HL) and no history of neurological disorder. The study was approved by the University of Maryland’s Institutional Review Board. All participants gave written informed consent and were compensated for their time.

Subjects came in on two different days for the MEG auditory task and structural MRIs. Neural magnetic signals were recorded in a dimly lit, magnetically shielded room with 160 axial gradiometer whole head MEG system (KIT, Kanazawa, Japan) at the Maryland Neuroimaging Center. The MEG data were sampled at 2 kHz, low pass filtered at 200 Hz and notch filtered at 60 Hz. Participants laid supine position during the MEG experiment while their head was in the helmet and as close as possible to the sensors. Each participant’s head shape was digitized using Polhemus 3SPACE FATRAK digitizer. The head position was tracked at the start and end of the experiment with 5 fiducial coils attached to their heads. Structural MRI data were collected on a Siemens 3T TIM Trio scanner with a 32-channel head coil. T1-weighted MP-RAGE images (magnetization-prepared rapid gradient-echo) were acquired in 208 sagittal slices with 0.8 mm isotropic voxels (256 × 256 matrix, TR = 2400 ms, TE = 2.01 ms, TI = 1060 ms, flip angle = 8◦, distance factor = 50%, GRAPPA (generalized auto-calibrating partially parallel acquisition) acceleration factor = 2, interleaved acquisition). MRI images were processed using recon-all pipeline in Freesurfer to generate the cortical reconstructions. MEG, MRI volumes and head shape digitization points were aligned using translation and rotation.

Subjects listened to 60 s-long audio segments (trial) from the audio book “The Legend of Sleepy Hollow”, by Washington Irving narrated by a male talker at three different background talker levels, speech in quiet, at 0 dB SNR and at −6 dB SNR (and a fourth condition, with a speech-babble background, that was not analyzed here). The background talker stimuli were 60 s-long audio segments from the same audio book narrated by a female talker. For the competing talker segments subjects were asked to attend to the foreground talker while ignoring the other. When mixing the two stimuli for the competing talker segments the sound level of the foreground talker was kept identical to the speech in quiet condition, while the sound level of the background talker was manipulated to change the noise level. Each condition included 3 trials.

During the task subjects were instructed to minimize the body movements as much as possible, and to keep their eyes open and fixate on a male/female cartoon face at the center of the screen (depending on which talker they were instructed to attend to). However, in this analysis only the trials of selectively attended to the male voice were analyzed, in order to limit computational processing time. Sound level was calibrated to approximately 70 dBA sound pressure level (SPL) using 500 Hz tones and equalized to be approximately flat from 40 Hz to 4 kHz. The stimuli were delivered with E-A-RTONE 3 A tubes (impedance 50 Ω), which strongly attenuate frequencies above 4 kHz, and E-A-RLINK (Etymotic Research, Elk Grove Village, United States) disposable earbuds inserted into the ear canals.

### Pre-processing and Data Cleaning

MNE-python 0.21.0 (Gramfort et al., 2013; Gramfort, 2014) has been used to perform all the pre-processing steps. Prior down sampling the recordings to 50 Hz, the noisy channels have been removed and temporal signal space separation (tsss) was used to eliminate the artifacts (Taulu & Simola, 2006). The data were filtered between 0.1 Hz and 25 Hz using a causal minimum phase FIR filter followed by applying independent component analysis to extract and remove cardiac and muscle artifacts (Makeig et al., 1995; Lee et al., 1999). After removing the initial 5 s of the data, the resulting 55 s were extracted and filtered to the desired frequency bands (0.1-4 Hz and 4-8 Hz) using causal minimum phase FIR filters.

### Granger Causality Analysis via NLGC

We use NLGC to extract causal interactions from the MEG recordings. NLGC models the underlying neural sources activities as sparse vector auto-regressive (VAR) processes and estimates the model parameters directly from the MEG recordings without explicitly solving the inverse problem (Soleimani et al., 2022). The forward model (Baillet et al., 2001) at a given time *t* is expressed as

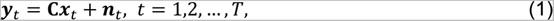

where ***y****_t_* ∈ ℝ^155^ denotes the MEG measurements vector, ***C*** ∈ ℝ^155×*M*^ shows the mapping between the *M* underlying sources and the MEG sensors which is also referred to as the lead-field matrix (Mosher et al., 1999), ***x****_t_* ∈ ℝ*^M^* indicates the *M* neural source activity vector, and ***n****_t_* ∈ ℝ^155^ represents the measurements noise vector which is typically considered as a zero-mean Gaussian process and its covariance can be estimated from the empty room recordings (Engemann & Gramfort, 2015). Given the fact that the sampling frequency is 50 Hz, the total number of time points in a 55-second trial is given by *T* = 50 × 55 = 2750.

The underlying VAR model representing the neural dynamics is given by

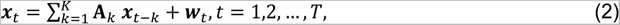

where **A***_k_* ∈ ℝ*^M^*^×*M*^ represents the coefficient matrix corresponding to the *k*^th^ lag (*k* = 1,2, …, *K*) and ***w****_t_* ∈ ℝ*^M^* is modeled as a zero-mean Gaussian process. It should be noted NLGC provides estimates of the VAR coefficients matrices and the noise covariance matrix along the way prior to the connectivity analysis. Here, the results are obtained with *K* = 2 lags which considers the history of the neural activities up to 40 ms (twice the sampling interval). Furthermore, to mitigate the computational cost of the analysis, the approximated version of the lead-field matrix ***C*** ∈ ℝ^155×*M*^ is used by summarizing the contribution of the neural sources dipoles in ‘ico – 1’ source-space with *M* = 84, using their first *r* principal components computed in ‘ico – 4’ source space with *M* = 5124 (Cheung et al., 2010; Babadi et al., 2014). As such, instead of considering all the original 5124 neural sources in the forward model, we proceed with *M* = *r* × 84. In other words, the brain is spanned with 84 non-overlapping regions as in ‘ico – 1’, however, to enhance the accuracy of the model, we utilize *r* principal components for each region calculated in ‘ico – 4’ source space. In this study, deploying *r* = 4 provides adequate resolution for the source space while significantly reducing the computation time. After determining the VAR model parameters, to identify the statistically significant GC links, NLGC performs a statistical test with false discovery rate (FDR) controlled at 0.1%,

NLGC evaluates the predictive influence of the past activity of one source on present activity of another over ‘ico-1’ source space and outputs an 84 × 84 connectivity matrix representing the directional connectivity across the dipoles. This matrix is then mapped to Desikan-Killiany atlas (Desikan et al., 2006) with the 68 anatomical regions of interest (ROIs). The considered ROIs for the connectivity analysis are as follows (Fig. S1):

- frontal: ‘rostralmiddlefrontal’, ‘caudalmiddlefrontal’, ‘parsopercularis’, ‘parstriangularis’.
- temporal: ‘superiortemporal’, ‘middletemporal’, ‘transversetemporal’.
- parietal: ‘inferiorparietal’, ‘posteriorcingulate’.

This way, each captured GC link can be assigned a source-target label as *S*_1_ → *S*_2_ where *S*_1_, *S*_2_ ∈ {frontal, temporal, parietal}. As a result, for each subject-condition-trial, the connectivity network in the auditory cortex can be summarized as 9-dimensional tuple where each entry shows the number of links corresponding to connectivity regions (e.g., frontal → temporal, parietal → parietal, etc.). It is worth mentioning that one can include hemispheres to identify inter- and intra-hemispheric connections as *S*_1_(*h*_1_) → *S*_2_(*h*_2_) with *S*_1_, *S*_2_ ∈ {frontal, temporal, parietal} and *h*_1_, *h*_2_ ∈ {Right, Left}. In this case, the representing tuple would have 36 entries. For more information on the parameter selection and model tuning, we refer the reader to (Soleimani et al., 2022).

### Average Power Calculations

To calculate the spatial distribution of average power across cortical sources, as shown in the gauges in Figs. 2, 3, 5, and 6, we utilize NLGC’s estimated time-courses of the cortical sources which are calculated in parallel with estimating the VAR model parameters. Further details can be found in (Soleimani et al., 2022).

Denoting by 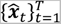 the estimated source activity time-courses, the average power of an ROI can be expressed as sum of the average power of all diploes lying within the ROI as:

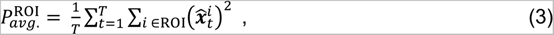

where 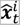 denotes the estimated activity of the *i*^th^ neural source.

### Determining Nature of Captured Links via NLGC

As mentioned earlier, NLGC can reliably estimates the VAR model parameters in Eq. (2). Given the estimated coefficients matrices 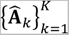, we can further categorize the links into three main classes. To elaborate, assume there is a link from the *j*^th^ to the *i*^th^ neural source. The resulting auto-regressive equation for source *i* can be written as

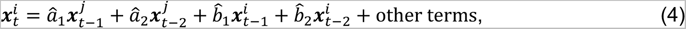

where 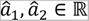 and 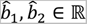 are the estimated cross-coupling and self-coupling coefficients, respectively, corresponding to the first and the second lags, respectively. It should be noted that since the *j* → *i* link exists, at least one of the coefficients is non-zero (Bressler & Seth, 2011). Depending on the sign of 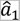 and 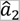, the nature of the link *j* → *i* can initially be categorized as follows:

- Facilitative: if 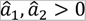, a boost in the activity of the *j*^th^ dipole would increase the *i*^th^ neural source activity within a delay of at most 40 ms.
- Suppressive: if 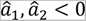, a reduction in the activity of the *j*^th^ dipole is followed by a decrement in the *i*^th^ neural source activity with a delay of at most 40 ms.

To identify the nature of the link for the other cases, i.e., 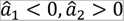 and 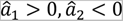, we consider the transfer function associated to the source-target pair (*j*, *i*) which is given by:

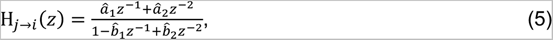

where 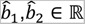 represent the estimated self-history coefficients corresponding to the first and second lags. Depending on the estimated coefficients (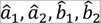), the transfer function behaves as a high- or low-pass filter. Interestingly, in this analysis, the shape of resulting filters for all captured GC interactions are determined only by the values of cross-coupling coefficients (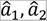) as the presence or absence of a link in the model is determined by these two coefficients. As such, we consider the nature of the links based on the resulting filter shape |H*_j_*_→*i*_(*z*)|. To account for potential numerical errors in estimation, we set a small margin ε = 0.01 and apply the following classification to determine the nature of the links, only if the absolute value of both estimated coefficients is greater than ε (See Fig. S2). Defining the binary variable 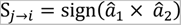, with

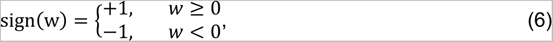

representing the signum function, we have, finally:

- Facilitative (analogous to excitatory): S*_j_*_→*i*_ > 0 with 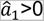 which results in a low-pass filter, i.e., an increase in the activity of the *j*^th^ ROI results in an increment in the overall activity of the neural source in the *i*^th^ ROI.
- Suppressive (analogous to inhibitory): S*_j_*_→*i*_ > 0 with 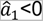 which also is a low-pass filter showing that a boost in the activity of the *j*^th^ ROI follows by a reduction in the overall activity of the *i*^th^ ROI.
- Sharpening (analogous to excitatory tuning plus lateral inhibition, except in time): S*_j_*_→*i*_ < 0 representing a high-pass filter. This case spans more complex situations which are neither facilitative nor suppressive, but rapid changes in sign (within a 40 ms window) in the activity of the *j*^th^ ROI affect the activity of the *i*^th^ ROI.

Fig. S2 shows a summary of the aforementioned link categories.

### Statistical Testing

All statistical analysis was performed in R (version 4.0.5) (R Core Team 2021). We used generalized linear mixed-effect models to analyze the relationship between GC link counts (dependent variable) and age (categorical variable; 2 levels: younger, older), condition (categorical variable; 3 levels: quiet, 0 dB, −6 dB), connectivity regions (categorical variable; 9 levels: T→T, T→F, …, P→P). For this we used glmmTMB (version 1.1.4) (Brooks et al., 2017) including truncated generalized Poisson distribution and adding zero-inflation for each frequency band. The best fit model out of the full model was selected by buildmer package (version 2.6) (Voeten, 2021).

To compare the effects of nature of the link type (categorical variable; 3 levels: facilitative, sharpening, suppressive), we modelled the data using Poisson log-normal distribution for each frequency band. In this model the dependent variable was GC link count, and the independent variables were age, condition, and the nature of the link type. For this analysis we aggregated the link counts over connectivity regions as connectivity regions was not a significant predictor. Based on a full model accounting for all the variables, the best fit model was selected by a two-step process of defining the maximal possible model followed by stepwise backwards elimination, implemented with the default settings in buildmer package. The effect size *d* was quantified as the ratio of the estimate for the fixed effect divided by the square root of the sum of variances of the random effects (Westfall et al., 2014).

For each model, model assumptions for overdispersion, heteroskedasticity and zero-inflation were examined and verified using the DHARMa package (version 0.4.5) (Hartig, 2021). The post-hoc differences among the levels of the effects were tested using pairwise comparisons based on estimated marginal means, with Holm corrections using the package emmeans (version 1.7.5) (Lenth et al., 2021). The summary of the statistical models is given in supplementary files (Table S1 and S2).

### Data and software availability

The MEG and MRI data will be made available upon acceptance.

A python implementation of NLGC is publicly available on GitHub (Soleimani, 2022) to ease the reproducibility in the community.

## ACKNOWLEDGEMENTS

This work was supported by the National Institute on Aging (NIA) P01-AG055365 (to JZS), the National Institute on Deafness and Other Communication Disorders (NIDCD) R01-DC019394 (to JZS, BB), and the National Science Foundation SMA 1734892 (to JZS, BB), OISE 2020624 & CCF 1552946 (to BB).

## DISCLAIMER

The identification of specific products or scientific instrumentation is considered an integral part of the scientific endeavor and does not constitute endorsement or implied endorsement on the part of the authors, DoD, or any component agency. The views expressed in this article are those of the authors and do not necessarily reflect the official policy of the Department of Defense or the U.S. Government.

## SUPPLEMENTARY FILES

**Figure S1.**
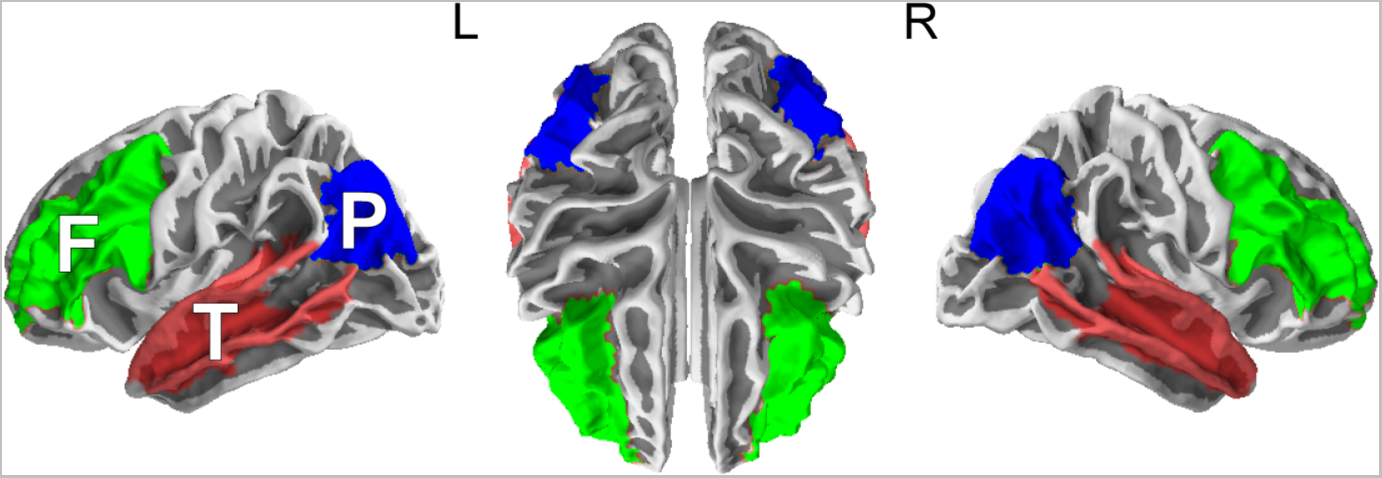
Auditory regions under investigation. From left to right, the studied auditory network on one of the participant’s brains is shown frontal (green), parietal (blue), and temporal (red) lobes.

**Figure S2.**
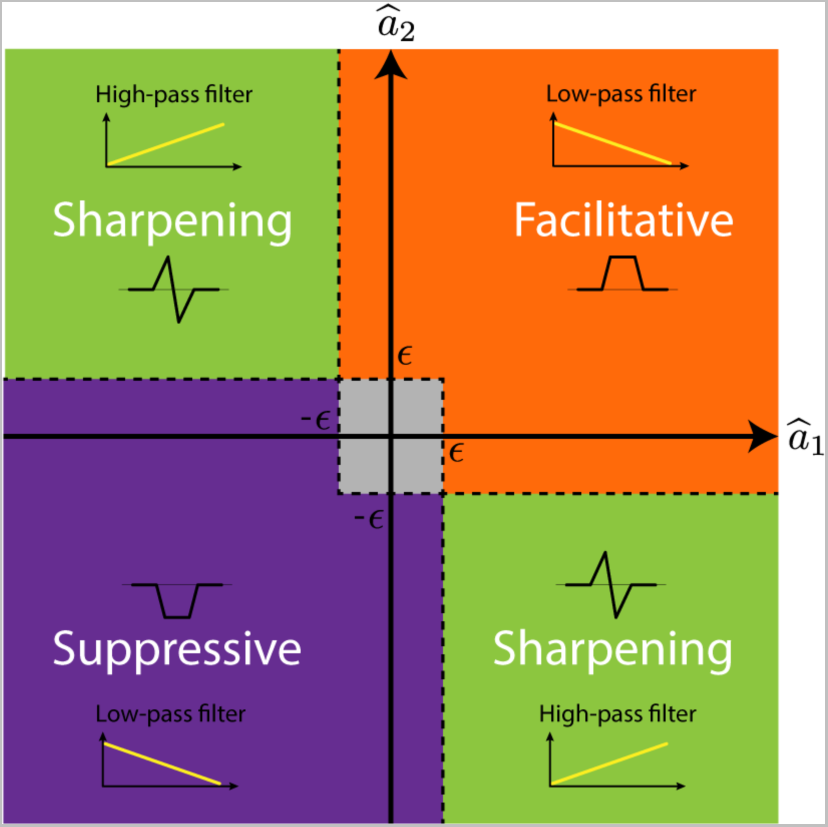
Classification rule based on cross-coupling coefficients for determining link nature. The figure illustrates how the relationship between the cross-coupling coefficients of a given target-source pair and the nature of the causal link. Facilitative and suppressive types are characterized by a low-pass filter, with positive coefficients for the former and negative coefficients for the latter. The sharpening, on the other hand, acts as a high-pass filter.

**Table S1.** Best-fitting statistical model (links ~ 1 + connected areas + age group + connected areas : age group + listening condition + Connected areas: listening condition + age group : listening condition + age group: listening condition : connected areas + (1| subject)) summary table corresponding to the connectivity analysis performed in Figs. 2,3,5, and 6.

|  | Delta Band |  |  | Theta Band |  |  |
| --- | --- | --- | --- | --- | --- | --- |
| Count Model | Estimate | Error | *p*-value | Estimate | Error | *p*-value |
| Intercept | -2.32 | 0.08 | $<2 \times 10^{-16}$ | -1.84 | 0.12 | $<2 \times 10^{-16}$ |
| F→P | -0.88 | 0.10 | $<2 \times 10^{-16}$ | -0.18 | 0.19 | 0.098 045 |
| F→T | -0.16 | 0.11 | 0.172 200 | -0.59 | 0.14 | 0.098 045 |
| P→F | -0.67 | 0.10 | $6.14 \times 10^{-12}$ | -0.26 | 0.15 | 0.000 149 |
| P→P | -1.36 | 0.09 | $<2 \times 10^{-16}$ | -0.12 | 0.16 | 0.323 995 |
| P→T | -0.47 | 0.14 | 0.006 532 | -1.23 | 0.13 | 0.828 220 |
| T→F | -0.57 | 0.10 | $7.26 \times 10^{-9}$ | -2.40 | 0.13 | $<2 \times 10^{-16}$ |
| T→P | -0.46 | 0.10 | $5.84 \times 10^{-6}$ | -0.40 | 0.15 | $<2 \times 10^{-16}$ |
| T→T | -0.86 | 0.17 | $1.74 \times 10^{-6}$ | -0.38 | 0.15 | 0.060 436 |
| Older | -1.23 | 0.10 | $<2 \times 10^{-16}$ | -1.06 | 0.14 | $1.11 \times 10^{-11}$ |
| 0dB | -0.23 | 0.11 | 0.026 244 | -0.08 | 0.16 | 0.960 726 |
| -6dB | -0.39 | 0.11 | 0.000 145 | -0.29 | 0.16 | 0.213 384 |
| F→P:Older | -1.47 | 0.13 | $<2 \times 10^{-16}$ | -0.24 | 0.21 | 0.072 143 |
| F→T:Older | -1.07 | 0.15 | $1.23 \times 10^{-13}$ | -0.05 | 0.17 | 0.382 401 |
| P→F:Older | -0.95 | 0.13 | $2.48 \times 10^{-14}$ | -0.34 | 0.19 | 0.221 654 |
| P→P:Older | -0.94 | 0.11 | $<2 \times 10^{-16}$ | -0.11 | 0.19 | 0.797 279 |
| P→T:Older | -1.09 | 0.19 | $2.64 \times 10^{-10}$ | -0.60 | 0.16 | 0.003 777 |
| T→F:Older | -2.05 | 0.16 | $<2 \times 10^{-16}$ | -2.39 | 0.16 | $<2 \times 10^{-16}$ |
| T→P:Older | -2.05 | 0.17 | $<2 \times 10^{-16}$ | -0.45 | 0.19 | 0.012 747 0 |
| T→T:Older | -0.82 | 0.22 | $3.44 \times 10^{-5}$ | -0.40 | 0.18 | 0.012 528 7 |
| F→P:0dB | -0.10 | 0.13 | 0.418 907 | -0.25 | 0.24 | 0.117 629 |
| F→T:0dB | -0.10 | 0.15 | 0.515 310 | -0.88 | 0.19 | $6.30 \times 10^{-7}$ |
| P→F:0dB | -0.04 | 0.13 | 0.515 310 | -0.02 | 0.21 | 0.957 421 |
| P→P:0dB | -0.32 | 0.12 | 0.006 586 | -0.64 | 0.21 | 0.000 284 |
| P→T:0dB | -0.04 | 0.19 | 0.580 661 | -0.33 | 0.19 | 0.137 191 |
| T→F:0dB | -0.49 | 0.14 | 0.000 902 | -2.28 | 0.19 | $<2 \times 10^{-16}$ |
| T→P:0dB | -0.85 | 0.16 | $1.74 \times 10^{-8}$ | -1.47 | 0.19 | $<2 \times 10^{-16}$ |
| T→T:0dB | -0.21 | 0.22 | 0.272 464 | -0.32 | 0.21 | 0.120 957 |
| F→P:-6dB | -0.11 | 0.12 | 0.406 904 | -0.03 | 0.24 | 0.862 018 |
| F→T:-6dB | -0.05 | 0.14 | 0.721 716 | -1.53 | 0.18 | $<2 \times 10^{-16}$ |
| P→F:-6dB | -0.36 | 0.13 | 0.005 262 | -0.07 | 0.20 | 0.146 881 |
| P→P:-6dB | -0.60 | 0.12 | $3.55 \times 10^{-7}$ | -0.04 | 0.21 | 0.498 756 |
| P→T:-6dB | -0.21 | 0.19 | 0.264 875 | -1.26 | 0.20 | $5.30 \times 10^{-10}$ |
| T→F:-6dB | -1.03 | 0.16 | $5.23 \times 10^{-10}$ | -2.23 | 0.18 | $<2 \times 10^{-16}$ |
| T→P:-6dB | -1.34 | 0.17 | $9.63 \times 10^{-15}$ | -0.53 | 0.19 | 0.003 35 |
| T→T:-6dB | -0.01 | 0.22 | 0.757 014 | -0.51 | 0.21 | 0.108 625 |
| Older:0dB | -0.59 | 0.14 | $9.24 \times 10^{-6}$ | -0.73 | 0.21 | 0.001 143 |
| Older:-6dB | -1.48 | 0.15 | $<2 \times 10^{-16}$ | -0.60 | 0.20 | 0.009 045 |
| F→P:Older:0dB | -0.37 | 0.18 | 0.041 652 | -0.04 | 0.30 | 0.563 227 |
| F→T:Older:0dB | -0.66 | 0.20 | 0.001 009 | -0.06 | 0.24 | 0.745 038 |
| P→F:Older:0dB | -0.65 | 0.17 | 0.000 140 | -0.53 | 0.27 | 0.061 382 |
| P→P:Older:0dB | -0.71 | 0.16 | $3.56 \times 10^{-6}$ | -0.28 | 0.27 | 0.432 418 |
| P→T:Older:0dB | -0.29 | 0.27 | 0.104 852 | -1.03 | 0.24 | $2.58 \times 10^{-5}$ |
| T→F:Older:0dB | -0.80 | 0.23 | 0.002 196 | -2.70 | 0.26 | $<2 \times 10^{-16}$ |
| T→P:Older:0dB | -1.31 | 0.12 | $7.64 \times 10^{-8}$ | -0.45 | 0.25 | 0.023 674 |
| T→T:Older:0dB | -0.04 | 0.30 | 0.961 451 | -0.74 | 0.27 | 0.006 775 |
| F→P:Older:-6dB | -0.99 | 0.19 | $6.13 \times 10^{-8}$ | -0.37 | 0.30 | 0.181 500 |
| F→T:Older:-6dB | -1.54 | 0.21 | $1.71 \times 10^{-14}$ | -1.16 | 0.23 | $2.30 \times 10^{-8}$ |
| P→F:Older:-6dB | -1.88 | 0.19 | $<2 \times 10^{-16}$ | -0.49 | 0.26 | 0.254 699 |
| P→P:Older:-6dB | -1.69 | 0.17 | $<2 \times 10^{-16}$ | -0.49 | 0.28 | 0.078 279 |
| P→T:Older:-6dB | -1.33 | 0.27 | $1.54 \times 10^{-6}$ | -0.95 | 0.25 | 0.000 412 |
| T→F:Older:-6dB | -1.92 | 0.25 | $8.31 \times 10^{-14}$ | -2.39 | 0.24 | $<2 \times 10^{-16}$ |
| T→P:Older:-6dB | -2.61 | 0.26 | $<2 \times 10^{-16}$ | -0.43 | 0.25 | 0.207 520 |
| T→T:Older:-6dB | -1.21 | 0.30 | $6.55 \times 10^{-6}$ | -1.6 | 0.26 | $5.69 \times 10^{-9}$ |
| AIC | 11 646.00 |  |  | 11 786.92 |  |  |
| Log Likelihood | 5764.00 |  |  | 5834.46 |  |  |
| Num. obs. | 1782 |  |  | 1782 |  |  |
| Random Effects |  |  |  |  |  |  |
|  | Variance | Std. Dev. |  | Variance | Std. Dev. |  |
| Intercept | 28 | 0.02417 |  | 0.1549 | 0.03936 |  |

**Table S2.**
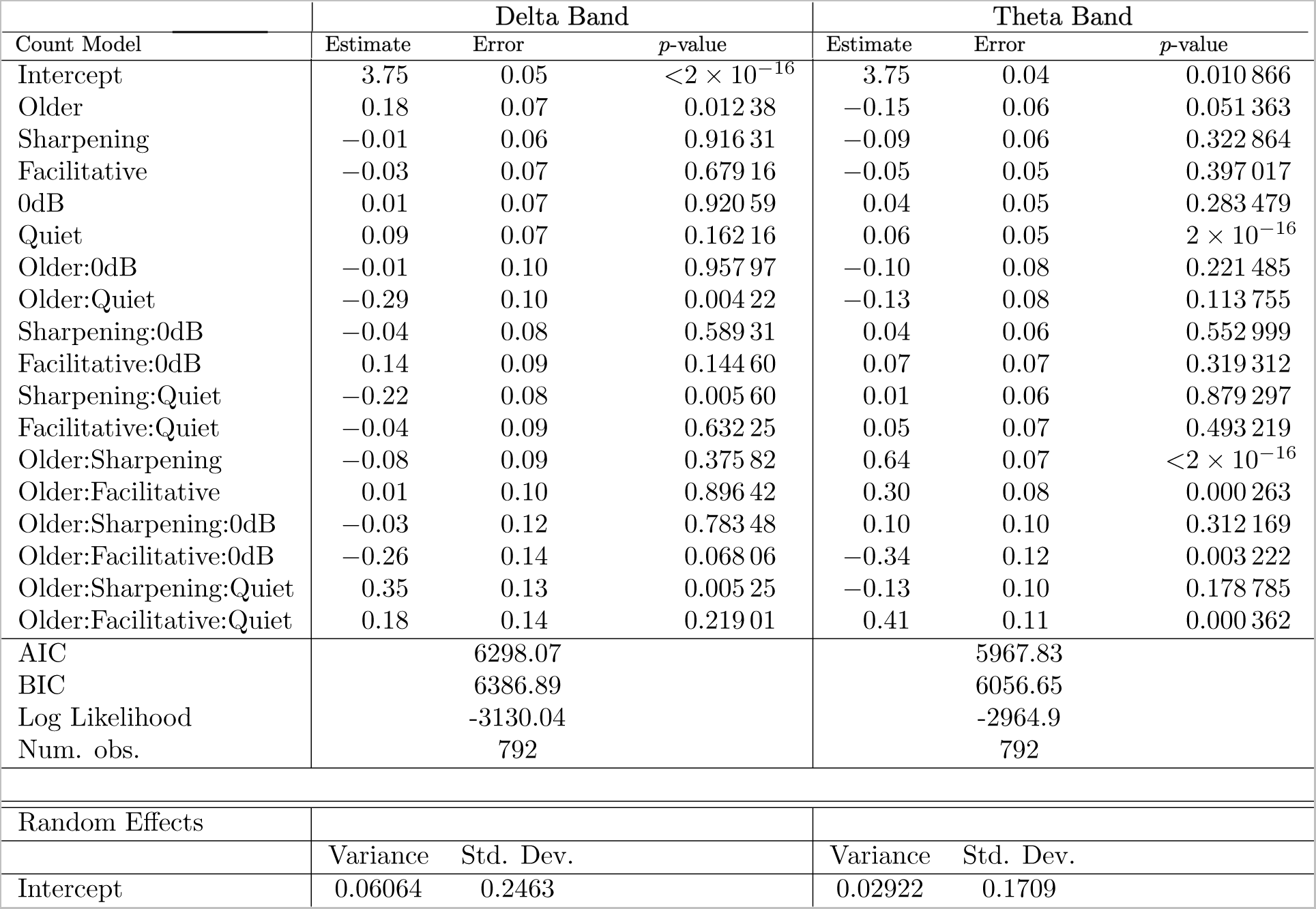
Best-fitting statistical model (links ~ 1+ age group + nature of links + listening condition + age group : listening condition + nature of links : listening condition + age group : nature of links + age group : nature of links : listening condition + (1|X), where X is observation-level random effect) summary table corresponding to determining nature of the links performed in Figs. 7 and 8.

## REFERENCES

Alain C, McDonald KL (2007) Age-related differences in neuromagnetic brain activity underlying concurrent sound perception, Journal of Neuroscience 27 1308–1314.

Anderson JS, Ferguson MA, Lopez-Larson M, Yurgelun-Todd D (2010) Topographic maps of multisensory attention, Proceedings of the National Academy of Sciences 107 20110–20114.

Anderson S, Parbery-Clark A, White-Schwoch T, Kraus N (2012) Aging affects neural precision of speech encoding, Journal of Neuroscience 32 14156–14164.

Babadi B, Obregon-Henao G, Lamus C, Matti H, Brown EN, Purdon PL (2014) A subspace pursuit-based iterative greedy hierarchical solution to the neuromagnetic inverse problem, NeuroImage 87 427–443.

Baillet S, Mosher J, Leahy RM (2001) Electromagnetic brain mapping, IEEE Signal processing magazine 14–30.

Bastiaansen MC, Van Der Linden M, Ter Keurs M, Dijkstra T, Hagoort P (2005) Theta responses are involved in lexical-Semantic retrieval during language processing, Journal of cognitive neuroscience 17 530–541.

Behroozmand R, Shebek R, Hansen DR, Oya H, Robin DA, Howard MA, Greenlee JDW (2015) Sensory-motor networks involved in speech production and motor control: An fMRI study, NeuroImage 109 418–428.

Binder JR, Desai RH, Graves WW, Conant LL (2009). Where is the semantic system? A critical review and meta-analysis of 120 functional neuroimaging studies, Cerebral Cortex, 19(12), 2767–2796.

Bressler S, Seth AK (2011) Wiener-Granger causality: a well established methodology, NeuroImage 58

Brooks ME, Kristensen K, Van Benthem KJ, Magnusson A, Berg CW, Nielsen A, Skaug HJ, Machler M, Bolker BM (2017) glmmTMB balances speed and flexibility among packages for zero-inflated generalized linear mixed modeling, The R journal 9 378–400.

Caspary DM, Milbrandt JC, Helfert RH (1995) Central auditory aging: GABA changes in the inferior colliculus, Experimental gerontology 30 349–360.

Caspary DM, Hughes LF, Ling LL (2013) Age-related GABAA receptor changes in rat auditory cortex, Neurobiology of aging 34 1486–1496.

Cheung BLP, Riedner BA, Tononi G, Van Veen BD (2010) Estimation of cortical connectivity from EEG using state-space models, IEEE Transactions on Biomedical engineering 57 2122–2134.

Desikan RS, Segonne F, Fischl B, Quinn BT, Dickerson BC, Blacker D, Buckner RL, Dale AM, Maguire PR, Hyman BT, Albert MS, Killiany RJ (2006) An automated labeling system for subdividing the human cerebral cortex on MRI scans into gyral based regions of interest, Neuroimage 31 968–980.

Dhamala M, Rangarajan G, Ding M (2008) Estimating Granger causality from Fourier and wavelet transforms of time series data, Physical review letters 100

Ding N, Simon JZ (2012) Emergence of neural encoding of auditory objects while listening to competing speakers, Proceedings of the National Academy of Sciences 109 11854–11859.

Dobri SGJ, Ross B (2021) Total GABA level in human auditory cortex is associated with speech-in-noise understanding in older age, Neuroimage 225 117474. DOI: 10.1016/j.neuroimage.2020.117474

Eckert MA, Cute SL, Vaden KI, Kuchinsky SE, Dubno JR (2012) Auditory cortex signs of age-related hearing loss, Journal of the Association for Research in Otolaryngology 13 703–713.

Engemann D, Gramfort A (2015) Automated model selection in covariance estimation and spatial whitening of MEG and EEG signals, NeuroImage 108 328–342.

Evans S, McGettigan C (2017) Comprehending auditory speech: previous and potential contributions of functional MRI, Language, Cognition and Neuroscience 32 829–846.

Fei N, Ge J, Wang Y, Gao J-H (2020) Aging-related differences in the cortical network subserving intelligible speech, Brain and Language 201

Friston KJ (2011) Functional and effective connectivity: a review, Brain connectivity 1 13–36. DOI: 0.1089/brain.2011.0008

Geerligs L, Renken RJ, Saliasi E, Maurits NM, Lorist MM (2015) A brain-wide study of age-related changes in functional connectivity, Cerebral cortex 25 1987–1999.

Gramfort A, Luessi M, Larson E, Engemann DA, Strohmeier D, Brodbeck C, Goj R (2013) MEG and EEG data analysis with MNE-Python, Frontiers in neuroscience 7

Gramfort A, Luessi, Martin, Larson, Eric, Engemann, Denis A., Strohmeier, Daniel, Brodbeck, Christian, Parkkonen, Lauri, Hamalainen, Matti S. (2014) MNE software for processing MEG and EEG data, Neuroimage 86 446–460.

Gross J, Kujala J, Hämäläinen M, Timmermann L, Schnitzler A, Salmelin R (2001) Dynamic imaging of coherent sources: studying neural interactions in the human brain, Proceedings of the National Academy of Sciences 98 694–699.

Harris KC, Dias JW, McClaskey CM, Rumschlag J, Prisciandaro J, Dubno JR (2022) Afferent loss, GABA, and Central Gain in older adults: Associations with speech recognition in noise, Journal of Neuroscience 42 7201–7212.

Hartig F (2021) DHARMa: residual diagnostics for hierarchical (multi-level/mixed) regression models,

Hipp JF, Engel AK, Siegel M (2011) Oscillatory synchronization in large-scale cortical networks predicts perception, Neuron 69 387–396.

Hughes LF, Turner JG, Parrish JL, Caspary DM (2010) Processing of broadband stimuli across A1 layers in young and aged rats, Hearing research 264 79–85.

Inui K, Nakagawa K, Nishihara M, Motomura E, Kakigi R (2016) Inhibition in the human auditory cortex, PLoS One 11

Karunathilake IMD, Dunlap JL, Perera J, Presacco A, Decruy L, Anderson S, Kuchinsky SE, Simon JZ (2023) Effects of Aging on Cortical Representations of Continuous Speech, J Neurophysiol DOI: 10.1152/jn.00356.2022

Klimesch W (1999) EEG alpha and theta oscillations reflect cognitive and memory performance: a review and analysis, Brain research reviews 29 169–195.

Kondo HM, Lin IF (2020) Excitation-inhibition balance and auditory multistable perception are correlated with autistic traits and schizotypy in a non-clinical population, Sci Rep 10 8171. PMCID: PMC7234986 DOI: 10.1038/s41598-020-65126-6

Kuchibhotla K, Bathellier B (2018) Neural encoding of sensory and behavioral complexity in the auditory cortex, Curr Opin Neurobiol 52 65–71. PMCID: PMC6924614 DOI: 10.1016/j.conb.2018.04.002

Kuchinsky SE, Vaden KI (2020) Aging, hearing loss, and listening effort: Imaging studies of the aging listener, Aging and Hearing: Causes and Consequences 231–256. DOI: 10.1007/978-3-030-49367-7_10

Lalwani P, Gagnon H, Cassady K, Simmonite M, Peltier S, Seidler RD, Taylor SF, Weissman DH, Polk TA (2019) Neural distinctiveness declines with age in auditory cortex and is associated with auditory GABA levels, Neuroimage 201 116033.

Lee T-W, Girolami M, Sejnowski TJ (1999) Independent component analysis using an extended infomax algorithm for mixed subgaussian and supergaussian sources, Neural computation 11 417–441.

Leicht G, Bjorklund J, Vauth S, Mu mann M, Haaf M, Steinmann S, Rauh J, Mulert C (2021) Gamma-band synchronisation in a frontotemporal auditory information processing network, Neuroimage 239

Lenth RV, Buerkner P, Gine-Vazquez I, Herve M, Jung M, Love J, Miguez F, Riebl H, Singmann H (2021) *emmeans: Estimated Marginal Means, aka Least-Squares Means*,

Mai G, Minett JW, Wang WSY (2016) Delta, theta, beta, and gamma brain oscillations index levels of auditory sentence processing, Neuroimage 133 516–528.

Makeig S, Bell A, Jung T-P, Sejnowski TJ (1995) *Independent component analysis of electroencephalographic data*,

Mosher J, Leahy R, Lewis PS (1999) EEG and MEG: forward solutions for inverse methods, IEEE Transactions on biomedical engineering 46 245–259.

Obleser J, Wise RJS, Dresner MA, Scott SK (2007) Functional integration across brain regions improves speech perception under adverse listening conditions, Journal of Neuroscience 27 2283–2289.

Osnes B, Hugdahl K, Specht K (2011) Effective connectivity analysis demonstrates involvement of premotor cortex during speech perception, Neuroimage 54 2437–2445.

Palva JM, Wang SH, Palva S, Zhigalov A, Monto S, Brookes MJ, Schoffelen J-M, Jerbi K (2018) Ghost interactions in MEG/EEG source space: A note of caution on inter-areal coupling measures, Neuroimage 173 632–643.

Palva S, Palva JM (2012) Discovering oscillatory interaction networks with M/EEG: challenges and breakthroughs, Trends in cognitive sciences 16 219–230.

Pantev C, Okamoto H, Ross B, Stoll W, Ciurlia-Guy E, Kakigi R, Kubo T (2004) Lateral inhibition and habituation of the human auditory cortex, European Journal of Neuroscience 19 2337–2344.

Pantev C, Okamoto, Hidehiko, Teismann, Henning (2012) Music-induced cortical plasticity and lateral inhibition in the human auditory cortex as foundations for tonal tinnitus treatment, Frontiers in systems neuroscience 6

Paraskevopoulos E, Chalas N, Bamidis P (2017) Functional connectivity of the cortical network supporting statistical learning in musicians and non-musicians: an MEG study, Scientific reports 7 1–10.

Paraskevopoulos E, Dobel C, Wollbrink A, Salvari V, Bamidis PD, Pantev C (2019) Maladaptive alterations of resting state cortical network in Tinnitus: A directed functional connectivity analysis of a larger MEG data set, Scientific Reports 9

Park H, Ince RAA, Schyns PG, Thut G, Gross J (2015) Frontal top-down signals increase coupling of auditory low-frequency oscillations to continuous speech in human listeners, Current Biology 25 1649–1653.

Parthasarathy A, Herrmann B, Bartlett EL (2019) Aging alters envelope representations of speech-like sounds in the inferior colliculus, Neurobiology of aging 73 30–40.

Peelle JE, Troiani V, Wingfield A, Grossman M (2010) Neural processing during older adults’ comprehension of spoken sentences: age differences in resource allocation and connectivity, Cerebral Cortex 20 773–782. DOI: 10.1093/cercor/bhp142

Peelle JE, Chandrasekaran, Keerthi, Powers, John, Smith, Edward E., Grossman, Murray (2013) Age-related vulnerability in the neural systems supporting semantic processing, Frontiers in aging neuroscience 5

Peelle JE, Wingfield, Arthur (2016) The neural consequences of age-related hearing loss, Trends in neurosciences 39 486–497.

Pu Y, Cheyne D, Sun Y, Johnson BW (2020) Theta oscillations support the interface between language and memory, NeuroImage 215

Ramamurthy DL, Recanzone GH (2020) Age-related changes in sound onset and offset intensity coding in auditory cortical fields A1 and CL of rhesus macaques, Journal of Neurophysiology 123 1015–1025.

Rauschecker JP, Scott SK (2009) Maps and streams in the auditory cortex: nonhuman primates illuminate human speech processing, Nature Neuroscience 718–724.

Richardson BD, Ling LL, Uteshev VV, Caspary DM (2013) Reduced GABAA receptor-mediated tonic inhibition in aged rat auditory thalamus, Journal of Neuroscience 33 1218–1227.

Riecke L, Esposito F, Bonte M, Formisano E (2009) Hearing illusory sounds in noise: the timing of sensory-perceptual transformations in auditory cortex, Neuron 64 550–561.

Ross B, Dobri S, Schumann A (2020) Speech-in-noise understanding in older age: The role of inhibitory cortical responses, European Journal of Neuroscience 51 891–908.

Rysop AU, Schmitt L-M, Obleser J, Hartwigsen G (2022) Age-related differences in the neural network interactions underlying the predictability gain, Cortex 154 269–286.

Sala-Llonch R, Bartrés-Faz D, Junqué C (2015) Reorganization of brain networks in aging: a review of functional connectivity studies, Frontiers in psychology 6 663. DOI: 10.3389/fpsyg.2015.00663

Schoffelen J-M, Gross J (2009) Source connectivity analysis with MEG and EEG, Human brain mapping 30 1857–1865.

Seth AK, Barrett A, Barnett L (2015) Granger causality analysis in neuroscience and neuroimaging, Journal of Neuroscience 35 3293–3297.

Seth AK, Edelman, Gerald M. (2007) Distinguishing causal interactions in neural populations, Neural computation 19 910–933.

Sharp DJ, Scott SK, Mehta MA, Wise RJ (2006) The neural correlates of declining performance with age: evidence for age-related changes in cognitive control, Cerebral Cortex 16 1739–1749.

Smith SM, Vidaurre D, Beckmann CF, Glasser MF, Jenkinson M, Miller KL, Nichols TE, Robinson EC, Salimi-Khorshidi G, Woolrich MW (2013) Functional connectomics from resting-state fMRI, Trends in cognitive sciences 17 666–682. DOI: 10.1016/j.tics.2013.09.016

Soleimani B, Das P, Karunathilake IMD, Kuchinsky SE, Simon JZ, Babadi B (2022) NLGC: Network localized Granger causality with application to MEG directional functional connectivity analysis, NeuroImage 260

Soleimani B, Das, Proloy (2022) *Network Localized Granger Causality MATLAB Implementation*. Available on GitHub: https://github.com/BabadiLab/NLGC

Stein A, Engell A, Okamoto H, Wollbrink A, Lau P, Wunderlich R, Rudack C, Pantev C (2013) Modulatory effects of spectral energy contrasts on lateral inhibition in the human auditory cortex: an MEG study, PLoS One 8

Tada M, Kirihara K, Koshiyama D, Fujioka M, Usui K, Uka T, Komatsu M, Kunii N, Araki T, Kasai K (2020) Gamma-band auditory steady-state response as a neurophysiological marker for excitation and inhibition balance: a review for understanding schizophrenia and other neuropsychiatric disorders, Clinical EEG and Neuroscience 51 234–243.

Tait L, Zhang J (2022) MEG cortical microstates: spatiotemporal characteristics, dynamic functional connectivity and stimulus-evoked responses, NeuroImage 251

Taulu S, Simola J (2006) Spatiotemporal signal space separation method for rejecting nearby interference in MEG measurements, Physics in Medicine & Biology 51

Tomasi D, Volkow ND (2012) Aging and functional brain networks, Molecular psychiatry 17 549–558. DOI: 10.1038/mp.2011.81

Upadhyay J, Silver A, Knaus TA, Lindgren KA, Ducros M, Kim D-S, Tager-Flusberg H (2008) Effective and structural connectivity in the human auditory cortex, Journal of Neuroscience 28 3341–3349.

Van Atteveldt N, Roebroeck A, Goebel R (2009) Interaction of speech and script in human auditory cortex: insights from neuro-imaging and effective connectivity, Hearing research 258 152–164.

Vanneste S, De Ridder D (2015) Stress-Related Functional Connectivity Changes Between Auditory Cortex and Cingulate in Tinnitus, Brain connectivity 5 371–383.

Voeten CC (2021) *buildmer: Stepwise Elimination and Term Reordering for Mixed-Effects Regression*,

Wang L, Saalmann YB, Pinsk MA, Arcaro MJ, Kastner S (2012) Electrophysiological low-frequency coherence and cross-frequency coupling contribute to BOLD connectivity, Neuron 1010–1020.

Westfall J, Kenny DA, Judd CM (2014) Statistical power and optimal design in experiments in which samples of participants respond to samples of stimuli, Journal of Experimental Psychology: General 143 2020. DOI: 10.1037/xge0000014

Willott JF, Parham K, Hunter KP (1991) Comparison of the auditory sensitivity of neurons in the cochlear nucleus and inferior colliculus of young and aging C57BL/6J and CBA/J mice, Hearing research 53 78–94.

Wingfield A, Grossman M (2006) Language and the aging brain: patterns of neural compensation revealed by functional brain imaging, Journal of neurophysiology 96 2830–2839. DOI: 10.1152/jn.00628.2006

Wong PC, Jin JX, Gunasekera GM, Abel R, Lee ER, Dhar S (2009) Aging and cortical mechanisms of speech perception in noise, Neuropsychologia 47 693–703. DOI: 10.1016/j.neuropsychologia.2008.11.032

Wong PCM, Ettlinger M, Sheppard JP, Gunasekera GM, Dhar S (2010) Neuroanatomical characteristics and speech perception in noise in older adults, Ear and hearing 31.

Wostmann M, Herrmann B, Maess B, Obleser J (2016) Spatiotemporal dynamics of auditory attention synchronize with speech, Proceedings of the National Academy of Sciences 113 3873–3878.

Zhang J, Cui Y, Deng L, He L, Zhang J, Zhang J, Zhou Q, Liu Q, Zhang Z (2016) Closely spaced MEG source localization and functional connectivity analysis using a new prewhitening invariance of noise space algorithm, Neural Plasticity

